# Computational Study of the Excitation of Human Induced Pluripotent Stem-Cell Derived Cardiomyocytes

**DOI:** 10.1101/2024.09.03.611054

**Authors:** Roshni Shetty, Raffi Samurkashian, Leslie Tung

## Abstract

Human induced pluripotent stem-cell derived cardiomyocytes (hiPSC-CMs) have proven to be a revolutionary advance for tissue engineering, disease modeling, and drug testing and discovery. Computational modeling enables a detailed electrophysiological analysis that is otherwise difficult or impossible to achieve under strictly experimental settings. Action potential characteristics of hiPSC-CMs measured in our lab at four different pacing rates were used it to modify the computational Kernik-Clancy hiPSC-CM model. The modified model was used to compare the excitation of single hiPSC-CMs with that of single human ventricular cardiomyocytes (hV-CMs) under varying conditions, including at stimulation at different strengths, rates and pulse durations. The physiological stimulation of both hiPSC-CMs and hV-CMs embedded within a tissue strand involves a biphasic waveform during which time excitatory currents (particularly *I*_Na_, but also *I*_CaT_ and *I*_CaL_ for hiPSC-CMs and *I*_NaL_ and *I*_CaL_ for hV-CMs) are activated during both phases of the waveform. *I*_Na_ in particular activated more slowly and with diminished amplitude under conditions of increasing pacing rate or increasing intracellular resistance. Lastly, histograms characterizing the relative amounts of excitatory currents in a population of hiPSC-CMs become broader with increasing levels of *I*_Na_ block, with *I*_CaT_ and *I*_CaL_ working in tandem to excite cells where *I*_Na_ has failed to activate. In general, hiPSC-CMs were found to be more excitable from rest compared with hV-CMs owing to their more depolarized resting potential and intrinsic automaticity despite a lower sodium channel density. Such a mismatch should be taken into consideration for applications using these cells, particularly for cardiac repair.

**Key Points:**

- Cardiomyocytes (CMs) generated from human stem cells derived from blood or skin have great potential for cardiac repair, safety pharmacology and disease modeling, but understanding their excitability is crucial for their proper application.
- Computational modeling reveals greater excitability of these cells in terms of various metrics compared with that of human adult ventricular CMs at multiple pacing rates.
- The excitation of CMs within a strand differs substantially from that usually used to study single isolated CMs and is dissected in terms of the underlying ionic currents.
- Computational modeling also predicts how a heterogeneous population of stem cell-derived CMs and their underlying ionic currents could respond to varying levels of reduced sodium current.
- Our study presents a cautionary note for applications using these cells, particularly for cardiac repair.

## Introduction

It has been several decades since human induced pluripotent stem cells (hiPSCs) have been differentiated into cardiomyocytes and used for cardiac research. These cells circumvent ethical issues relating to human tissue experiments and open the door to a diverse range of applications which include engineered tissues for cardiac repair, safety pharmacology, and disease modeling. In terms of cardiac repair, several Phase I and Phase II clinical trials are ongoing using hiPSC-CMs to treat various forms of cardiovascular disease (Sridharan *et al*., 2023), although pre-clinical studies implanting human embryonic stem cells into guinea pig and into non-human primate hearts have shown a transient period of tachyarrhythmias following cell implantation (Shiba *et al*., 2016; Romagnuolo *et al*., 2019; Cheng *et al*., 2023). For cardiotoxicity screening, the Comprehensive in vitro Proarrhythmic Assay (CiPA) initiative (Fermini *et al*., 2016; Colatsky *et al*., 2016) has been proposed to be a new standard for safety pharmacological testing, in which hiPSC-CMs are incorporated as a test platform. For disease modelling, hiPSC-CMs bearing mutations linked to specific heritable cardiac diseases are investigated for their arrhythmia mechanisms. A common thread among these applications is the specter of arrhythmias, which are either problematic (in the case of cardiac repair) or are a key metric (in the cases of cardiotoxicity screens and disease models).

Electrophysiological assays of hiPSC-CM have focused largely on prolongation of their action potentials (APs), formation of early afterdepolarizations, and irregular beat rate. Less attention has been given, however, to a fundamental property of these cells – the biophysics underlying their excitation. A cell that is unable to generate an action potential will be unable to conduct an electrical impulse. In tissue, this can introduce conduction block, which in turn may result in a break in the spread of electrical activity that subsequently leads to wavefront curvature and potentially, reentrant waves. The frequency of beating is an important parameter in this regard, because excitation of the cell is rate-dependent. At elevated beat rates, a population of cells with slightly different electrophysiological properties can develop variably sized subpopulations that either excite or fail to excite, particularly when challenged by different degrees of block of an underlying excitatory current.

In this study, the property of cellular excitation at the threshold of action potential generation is examined in detail using a computational model of the hiPSC-CM, and because these cells have been proposed to be surrogates for adult human cardiomyocytes, we also performed parallel studies on a model of the adult human ventricular cardiomyocyte for comparison. This allows the precise study of electrophysiological events which are otherwise challenging to observe experimentally and provides valuable insight into the underlying biophysics. Because such models are generally defined from diverse data sets from multiple laboratories, we constrained our hiPSC-CM model according to experimental measurements from our laboratory. Specifically, we placed special emphasis on action potential characteristics and ensured that our in-silico model reiterates these characteristics under paced conditions, not only at a single rate but at multiple rates.

The model is then used to investigate cellular excitation in terms of stimulus threshold and the strength-duration relation. We also compare the processes of excitation of cells in isolation vs. cells coupled together as a one-dimensional strand to delineate the consequences of electrical load. Finally, we examine the excitation of a heterogeneous population of hiPSC-CMs with reduced levels of sodium channel current.

## Methods

### 1. Single cell model of hiPSC-CM

Several computational models of the cellular electrophysiology of the hiPSC-CM have been published in recent years – Paci-Entcheva (Paci *et al*., 2020), Koivumaki-Tavi (Koivumäki *et al*., 2018), and Kernik-Clancy (Kernik *et al*., 2019). The source codes for the Paci-Entcheva and Kernik-Clancy models were readily available online so, we performed preliminary simulations of the APs generated by these models at different pacing rates. With the Paci-Entcheva model, the duration of the AP did not decrease monotonically at shorter pacing cycle lengths (CLs) as is observed experimentally, and simulations had a longer runtime compared with the Kernik-Clancy model.

For these reasons, we based our study on a modified version of the Kernik-Clancy (mKC) model. Because this model represents the electrophysiology of hiPSC-CMs, which are widely known to have an immature phenotype (Goversen *et al*., 2018), we also conducted parallel simulations of an adult human ventricular cardiomyocyte (hV-CM) with endocardial phenotype for comparison, using the Tomek-Rodriguez revision of the dynamic O’Hara-Rudy (ToR-ORd) biophysical model (Tomek *et al*., 2019). Compared with the ToR-ORd model, the mKC model has lower expression levels of several ionic currents (including *I*_Na_, *I*_K1_), a less negative diastolic transmembrane potential, and spontaneous activity typically found in hiPSC-CMs.

### 2. Syncytial model: linear strand

Linear strands of cardiac tissue were created in the form of one-dimensional cables of electrically coupled hiPSC-CMs. The geometric form and parameters of the strand are shown in **Fig. 1**. *R*_i_ is the intracellular axial resistance per cell (equal to the sum of myoplasmic and gap junctional resistances per cell), and *C*_m_ is the membrane capacitance.

**Figure 1.**
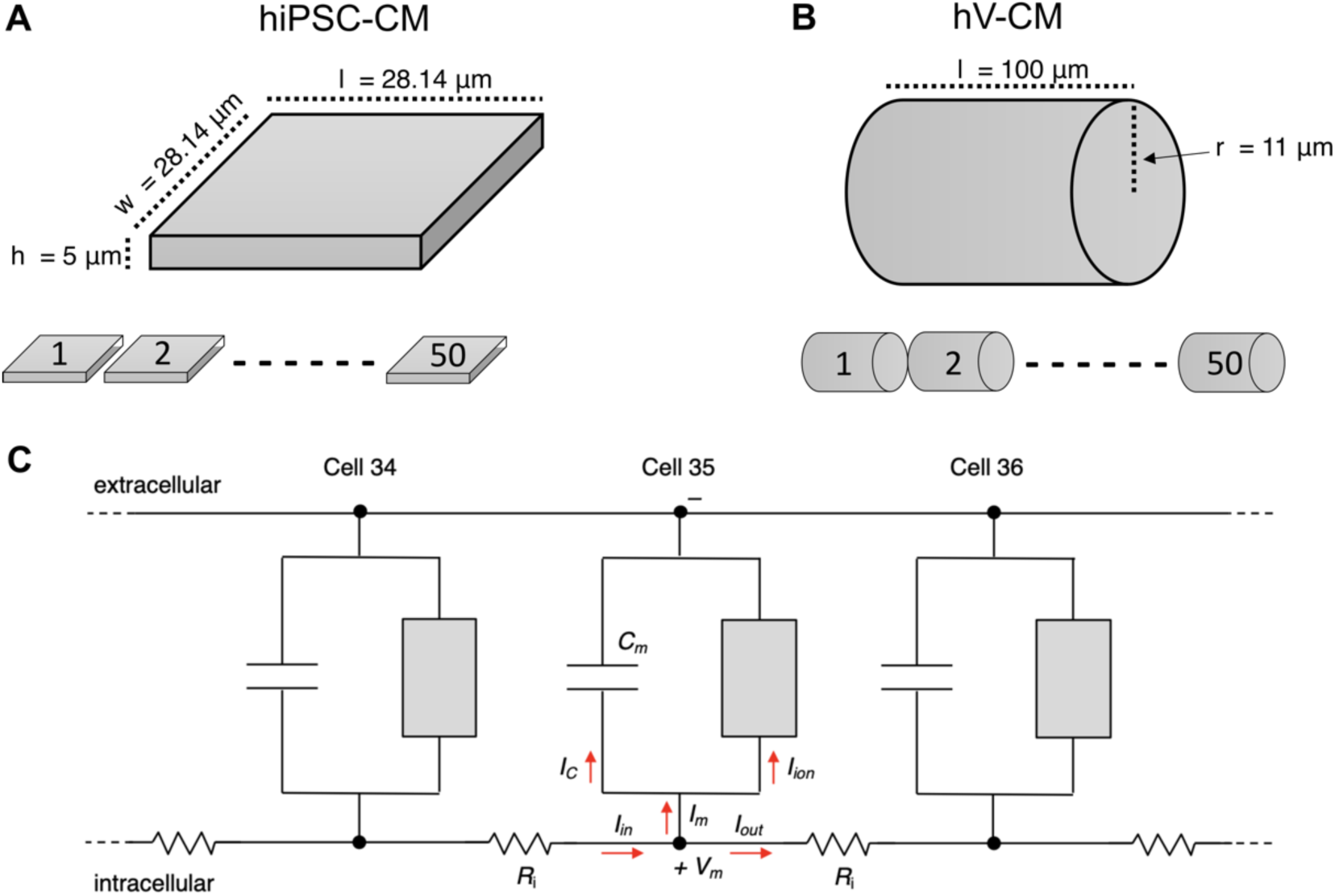
Geometric parameters for single cell and strand models. (**A**) hiPSC-CM dimensions and arrangement along a strand 50 cells in length. (**B**) hV-CM dimensions and arrangement along a strand with similar cell length. **C.** Electrical model of the middle of strand, showing cells 34, 35 and 36. Gray rectangle represents the cellular ion channels and electrogenic exchangers and pumps. *V*_m_ is transmembrane voltage, *I*C is capacitive current; *I*ion is the sum of currents from ion channels, exchangers and pumps; *I*m is total membrane current; *I*in is ionic current flowing into the cell; and *I*out is current flowing out of the cell. *R*_i_ is the intracellular axial resistance per unit length.

The currents in **Fig. 1C** have the following relationships:

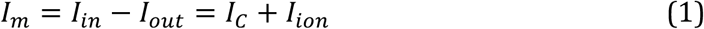

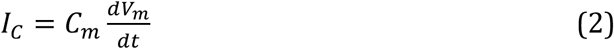

The original KC model for the hiPSC-CM specified an average cell volume of 3960 μm^3^ and cell capacitance of 60 pF. Although hiPSC-CMs in culture have complex shapes (Hwang *et al*., 2015), for simplicity we approximated the hiPSC-CM as a square tile with length and width of 28.14 μm and a height of 5 μm to match the volume defined in the model. For the hV-CM, we approximated the cell as a cylinder with length of 100µm, radius of 11µm, and capacitance of ∼76.7 pF (assuming an area-specific capacitance of 1µF/cm^2^) as specified in the ToR-ORd model. The capacitance of the mKC cell is calculated to be 60pF, which may be a bit high relative to the ToR-ORd cell capacitance, considering the smaller size and shortage of T-tubule development in hiPSC-CMs compared with hV-CMs. Also, the area-specific capacitance of the mKC cell becomes 2.80 µF/cm^2^ for the tile shape (or higher for more cylindrical or spherical cell shapes), as opposed to 1 µF/cm^2^ for the ToR-ORd cell. However, we chose not to modify this value and retained it from the original model formulation.

Both strands were constructed with 50 cells lined up end-to-end (**Fig. 1**). The first five cells of the strand were stimulated, and excitation was studied at the beginning and middle of the strand. Propagation was characterized at the middle of the strand.

### 3. Stimulation protocol

In general, when measuring the stimulus threshold for cells or strands with fixed *R*_i_, a standard conditioning train of 100 stimulus pulses (5 ms duration) was applied at a basic CL of 1000 ms, with a stimulus strength set to twice the minimum value needed for 1:1 capture for the 100 beats, after which a test pulse was applied with varying strength or duration at the 101st beat. For the excitation of cells and strands paced with variable CL, the models were run with standard duration pulses and the test CL during all 101 beats. For the excitation of strands with variable *R*_i_, the new value of *R*_i_ was applied during the standard conditioning train and test pulse. For the sensitivity analysis of strength-duration parameters to ion channel currents, the standard conditioning train of stimulus pulses was applied, after which the ion channel conductance was perturbed during the 101st beat. For the population simulations, a conditioning train of 100 stimulus pulses was applied at a variable CL for each model using stimulus strengths of 1.2x, 1.5x, or 2.0x the stimulus threshold specific to that model and pacing rate, after which *I*_Na_ conductance was decreased to simulate sodium channel blockade for another 10 beats.

### 4. Software

All simulations were performed in MATLAB (MathWorks, Natick, MA, USA). The ordinary differential equations (ODEs) in the single-cell models were solved using the MATLAB ode15s function. For the cable ODEs governing voltage and concentrations, we employed the forward-Euler method, while the gating variables were updated using the Rush-Larsen method. All curve fits were conducted with the MATLAB Curve Fitting Toolbox utilizing the trust region algorithm and the Nonlinear Least Squares method.

## Results

### 1. Modified Kernik-Clancy (mKC) hiPSC-CM model

The original KC model of the hiPSC-CM is based on data from multiple labs. Because the experimental conditions such as cell line, days of differentiation, cell density, and bath composition vary from lab to lab, we collected optical mapping data from our lab obtained from hiPSC-CM monolayers derived from the WTC11 cell line (Miyaoka *et al*., 2014) at 28-30 days of differentiation and at plating densities of 300,000±10% cells/cm^2^. We measured action potential duration at 80% repolarization (APD_80_) and action potential duration at 30% repolarization (APD_30_) at four different pacing rates (2000, 1000, 700 and 500 ms CL – corresponding to 0.5, 1.0, 1.43 and 2.0 Hz, respectively) to determine the range and variability of these parameters. Although these data were collected from multicellular and not single cell preparations, we subsequently confirmed in our simulations that APD_80_ and APD_30_ of the propagating AP in the strand had less than 1.5% root mean square difference (RMSD) from those of the single cell stimulated at twice threshold at the four pacing rates. **Fig. 2A** illustrates the variation of APD_80_ measured across a single monolayer at a pacing CL of 1000 ms.

**Figure 2.**
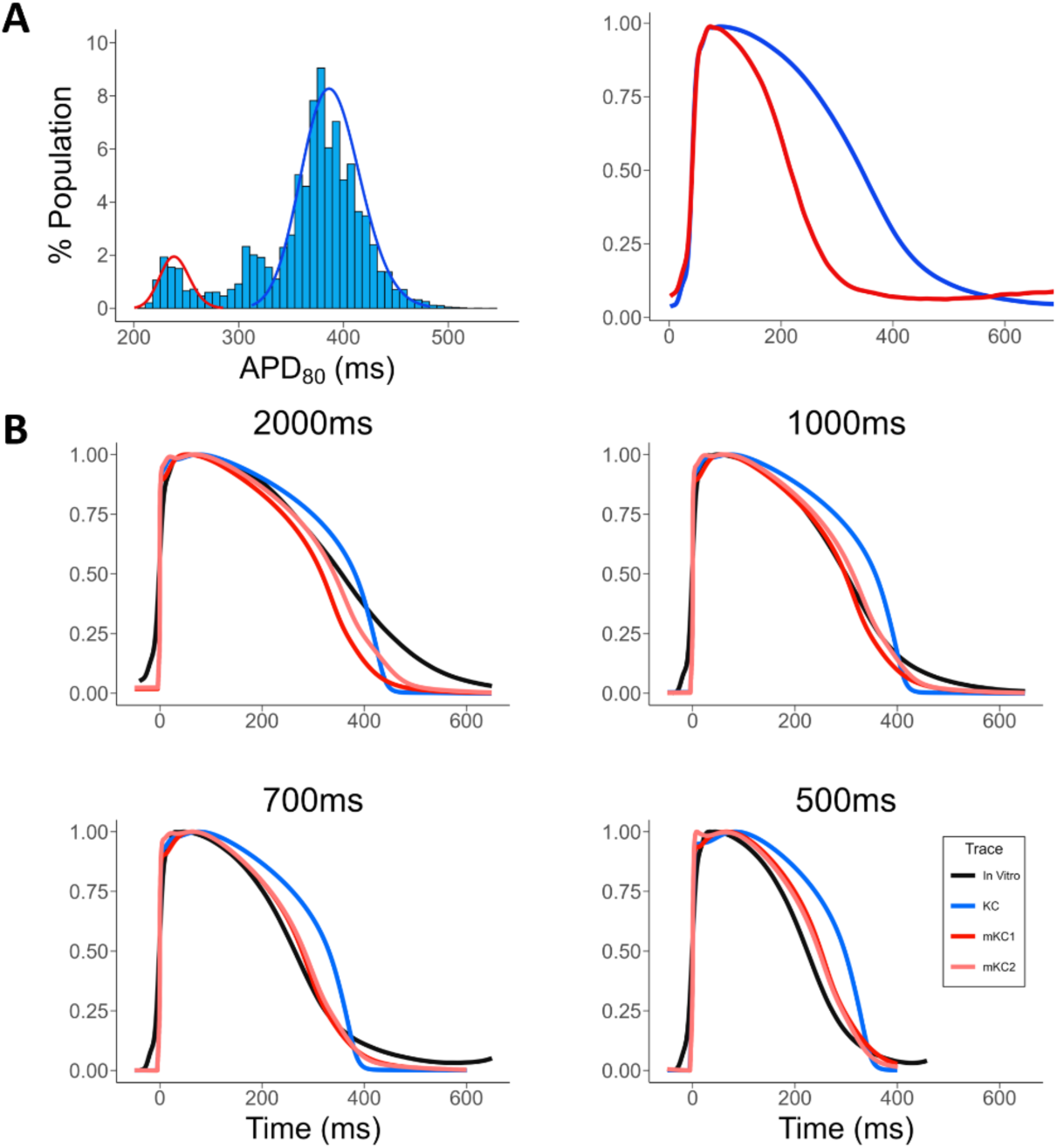
Experimental and computational hiPSC-CM APs. (**A**) Histogram (top left) of APD80 values at 1Hz pacing collected from 31 samples across 6 batches of WTC11 iPSC-derived cardiomyocytes. Red and blue Gaussian curves denote the presence of two distinct populations of cardiomyocytes. Red and blue APs (top right) reconstructed from optical mapping recordings corresponding to the distinct populations of cardiomyocytes shown in the histogram. (**B**) Comparison of in vitro (black), original KC (blue), mKC1 (red) and mKC2 (orange) APs at 2000ms (middle left), 1000ms (middle right), 700ms (bottom left), and 500ms (bottom right) pacing cycle lengths. In vitro APs at each pacing rate (black traces) are shown, averaged across the population of APs with larger mean APD80 (corresponding to the blue group in **A**). AP traces have been normalized from 0 to 1 to mimic the optical measurements of APs.

Kernel distribution fitting of the data suggests at least two subpopulations of cells, or perhaps three at some pacing rates, with significantly different mean APD_80_ values, which might reflect different emerging phenotypes. The averaged APs of the two subpopulations are also shown in **Fig. 2A**. In general, we fit the histogram of each monolayer sample with two or three Gaussian distributions and used only the distribution with the largest mean for this study. The averaged APs of that distribution are shown in **Fig. 2B** (black traces) for each of the four pacing rates. The upper and lower 10% values of the distribution were entered into **Table 1** as a measure of the range of APD_80_ values. This procedure was repeated for the other three pacing rates and then carried out again for APD_30_ and the ratio APD_30_/APD_80_ (as a measure of AP shape) at all four pacing rates. Experimentally measured values of conduction velocity (*θ*) are also listed.

**Table 1.**
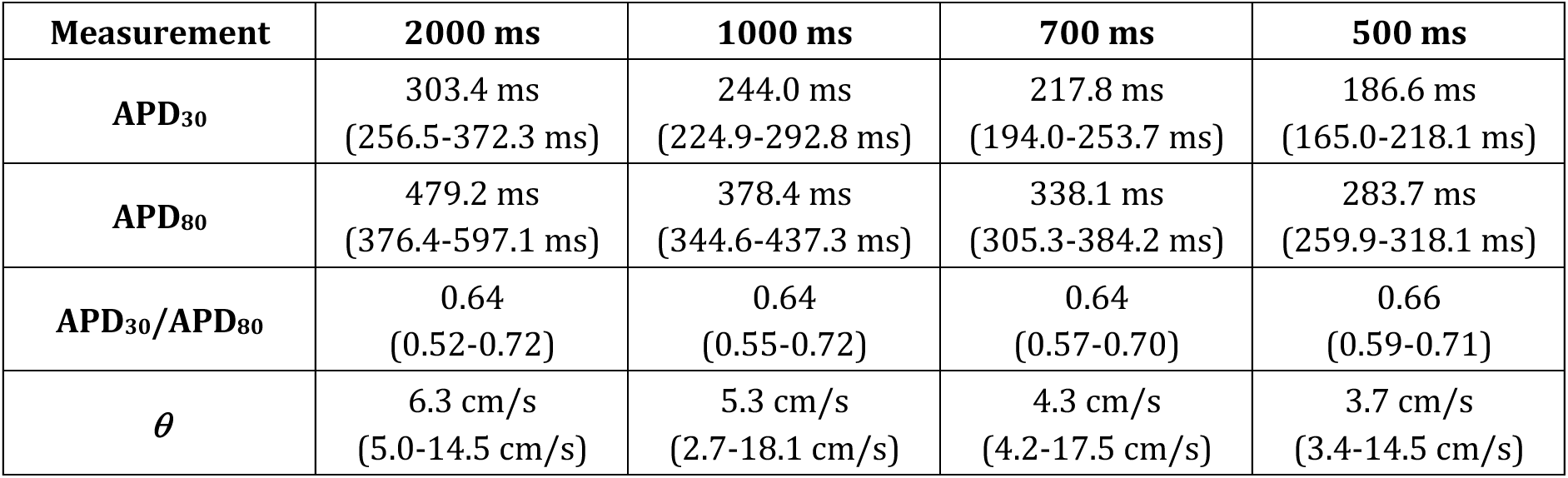
In vitro electrophysiological constraints used for population modeling. Constraints on AP parameters were derived from *in vitro* data as the upper and lower 10% cutoffs of Gaussian curves fitted to APD30, APD80, and APD30/APD80 histograms. Measurements of *θ* are also listed. Data are presented as mean (spread).

| Measurement | 2000 ms | 1000 ms | 700 ms | 500 ms |
| --- | --- | --- | --- | --- |
| APD<sub>30</sub> | 303.4 ms<br>(256.5-372.3 ms) | 244.0 ms<br>(224.9-292.8 ms) | 217.8 ms<br>(194.0-253.7 ms) | 186.6 ms<br>(165.0-218.1 ms) |
| APD<sub>80</sub> | 479.2 ms<br>(376.4-597.1 ms) | 378.4 ms<br>(344.6-437.3 ms) | 338.1 ms<br>(305.3-384.2 ms) | 283.7 ms<br>(259.9-318.1 ms) |
| APD<sub>30</sub>/APD<sub>80</sub> | 0.64<br>(0.52-0.72) | 0.64<br>(0.55-0.72) | 0.64<br>(0.57-0.70) | 0.66<br>(0.59-0.71) |
| $\theta$ | 6.3 cm/s<br>(5.0-14.5 cm/s) | 5.3 cm/s<br>(2.7-18.1 cm/s) | 4.3 cm/s<br>(4.2-17.5 cm/s) | 3.7 cm/s<br>(3.4-14.5 cm/s) |

After adjusting extracellular sodium concentration from 140 mM to 135 mM to match our experimental conditions, we found that while the KC model could be pace-captured (entrained) at the 1000 ms (1 Hz), 700 ms (1.43 Hz) and 500 ms (2 Hz) pacing CLs, it could not be pace-captured for one out of six batches of hiPSC-CMs at the 2000 ms (0.5 Hz) CL.

Increasing *g*_K1_ is known to provide a more stable resting potential and is thought to be a sign of maturation (Vaidyanathan *et al*., 2016). With *g*_K1_ at twice its original value, the KC model could be paced at all CLs, but APD_30_, APD_80_ and APD_30_/APD_80_ of the KC model failed to fall within the experimental range at 500 ms when the constraints at 2000 ms CLs were satisfied, or at 2000ms when the constraints at 500 ms were satisfied.

We also observed a significant difference in AP morphology – our in vitro APs had a more prolonged tail of repolarization and a more triangular shape than that of the KC model (**Fig. 2B**), as has been reported by others (Paci *et al*., 2013; Crestani *et al*., 2020; Gunawan *et al*., 2021; Seibertz *et al*., 2023). Experimentally, APD_30_/APD_80_ (a measure of shape) was consistently smaller across the four pacing rates (ranging from 0.52 to 0.72 with mean between 0.64 and 0.66) than that of the KC model (around 0.76). We were not able to reproduce the tail in the KC model by varying only ion channel conductances, permeabilities and transporter rates. The only simple adjustment that we could identify which could prolong the tail was an increase in the *α* component (rate constant from closed to open state) of the *I*_Kr_ activation gating parameter, which effectively scaled down its time constant, so *α* was increased by a factor of 30. It was also necessary to decrease the permeability, *p*_CaL_, of the L-type calcium channel by 55% to shorten the AP into the experimental range, and to decrease *I_f_* (by 50%) and slightly increase *g*_K1_ (by 10%) so that the model could be paced at 0.5 and 1 Hz while still exhibiting stable spontaneous activity. The exact values of these changes are summarized in **Table 2** as model 1 (mKC1), which we consider to be a minimal modification of the KC model. This model replicates AP shape very well at 1Hz (**Fig. 2B**) and is robust for small variations in the model conductances but fits our pacing rate data at 0.5 and 2.0 Hz only at the limits of their experimental ranges. The ratio of APD_30_/APD_80_ varied from 0.68 to 0.69 across the four pacing rates – a better oit to our experimental data than that of the original KC model.

**Table 2.** Parameters for the hiPSC-CM models.

| Model 1 (mKC1) | Model 2 (mKC2) |
| --- | --- |
| Xr1_1: 30 times original value | Xr1: 16.1404 times original value |
| gK1: 1.1 times original value | gK1: 2 times original value |
| p_CaL: 0.45 times original value | tauf_const: 0.8090 times original value |
| gf: 0.5 times original value | Xs_voltage shift: 44.518 mV left shift |
| | g_PCa: $0.995 \times 10^{-3}$ times original value |

We also found a second set of parameters which provided a better match to the rate-dependent changes in APD, but this involved altering a larger number of parameters (see **Appendix** for methodology). This version is summarized in **Table 2** as model 2 (mKC2), and its APs are displayed in **Fig. 2B**. Here, the *α* component of the *I*_Kr_ activation gating parameter *Xr1* was scaled up by a factor of about 16, SL pump activity was reduced by a factor of 1000, *I*_Ks_ activation was shifted negatively by 44 mV, and *I*_CaL_ voltage-dependent inactivation gate *τ* was reduced by 18%. In addition, *I*_K1_ conductance was doubled to allow pacing at 1 and 0.5 Hz, similar to its increase in the KC mature sub-model (Kernik *et al*., 2019). The mKC2 model matches AP shape adequately and more closely replicates the rate-dependent changes in APD and ratio of APD_30_/APD_80_ (0.68 across all four pacing rates) of our *in vitro* data (**Fig. 17**). Nevertheless, we were unable to find a set of parameters yielding a modified KC model that would oit all our experimental data at the center of their measured ranges. More specifically, the slope of the rate-dependence of APD_80_ on pacing rate was never as high as that measured experimentally. We found this discrepancy to be a common shortcoming of the all the variant models that we tested (**Appendix Fig. A1**). In general, results in this study for the mKC1 and mKC2 models were very similar except where noted, while heterogeneous population models using mKC1 had less scatter than those using mKC2. Hence, results mainly using mKC1 cells are presented here.

### 2. Stimulus threshold (*I*_T_)

#### 2.1. Stimulus threshold for the single cell

For the mKC1 single cell, initial conditions as specified in the original KC model code (https://github.com/ClancyLabUCD/IPSC-model) were used, and the parameters were modified according to **Table 2**. For the ToR-ORd single cell, initial conditions specified in the model code were used (https://github.com/jtmff/torord).

The cell was stimulated by a rectangular current pulse applied intracellularly and traversing the cell membrane. Simulations began for 100 beats at a pacing cycle length (CL) of 1000 ms, a pulse duration (*d*) of 5 ms, and a stimulus strength set to twice the minimum value needed for 1:1 capture for the 100 beats. Following this conditioning interval, the stimulus strength was varied for the 101^st^ beat (with *d* kept at 5 ms) to determine the stimulus threshold (*I*_T_) for that beat, which we defined to be the minimum strength required to elicit an AP with a peak in transmembrane potential (*V*_m_) ≥ 0 mV.

Both the hiPSC-CM and hV-CM show an all-or-none response (**Fig. 3**), with *I*_T_ of 1.49 pA/pF for the hiPSC-CM and 5.83 μA/μF for the hV-CM. *V*_m_ exhibits a threshold potential (*V*_T_), which is –69.1 mV for the hiPSC-CM and –63.6 mV for the hV-CM. In contrast, for the original 2011 ORd model of the hV-CM, the voltage response is graded, with a *V*_T_ that is not sharply defined but roughly around –33 mV. Also, the subthreshold response of the AP displays some active behavior, as indicated by a relatively long interval of elevated potential before return to rest (**Appendix Fig. A2**).

**Figure 3.**
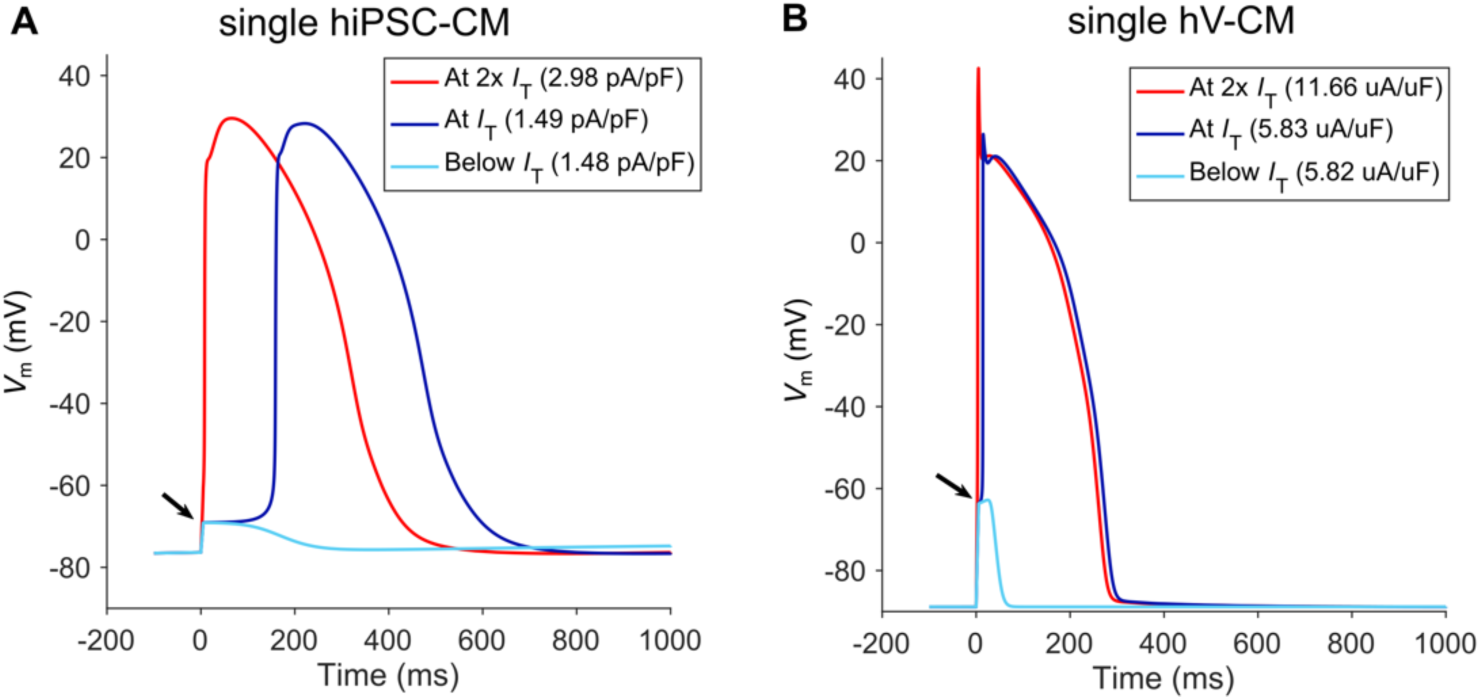
Single cell excitation at different stimulus strengths. (**A**) hiPSC-CM (mKC1 model). (**B**) hV-CM (ToR-ORd model). Cells were conditioned by a stimulus train of 100 pulses (1000 ms CL) after which they were stimulated 1 s later with a current pulse having strength either just below threshold (cyan trace), at threshold (*I*T, blue trace), or at twice threshold (red trace). Shown are the transmembrane potential (*V*m) responses. The black arrow indicates the threshold potential (*V*T).

A feature that is prevalent at threshold in the single cell models but is experimentally difficult to capture consistently is the lag time (latency) for activation (or time from stimulus onset to maximum upstroke, *t*_lag_) of the AP, a parameter that is seldom reported but has been related to excitability (Davidenko *et al*., 1990). For the given stimulus conditions, the hiPSC-CM has a long *t*_lag_ of ∼159 ms, and hV-CM has a much shorter *t*_lag_ of ∼14 ms (**Fig. 3**), while the ORd hV-CM has a *t*_lag_ of ∼187 ms (**Appendix Fig. A2**). Note, however, that the exact value of *t*_lag_ depends on the resolution of step sizes in stimulus strength, which was 0.01 pA/pF for hiPSC-CM and 0.01 µA/µF for hV-CM when normalized to cell capacitance. As the stimulus strength is increased above the stimulus threshold, *t*_lag_ becomes very short. At twice threshold, *t*_lag_ is ∼8 ms in hiPSC-CMs and ∼3 ms in hV-CMs. Below threshold, *V*_m_ returns to a constant negative resting potential for hV-CMs, but for hiPSC-CMs, it reaches a maximum diastolic potential and then slowly depolarizes due to the intrinsic ability of the cell to fire in the absence of electrical stimulation.

The ionic currents underlying the AP vary in terms of the voltage range and time interval when they are most active (**Fig. 4**).

**Figure 4.**
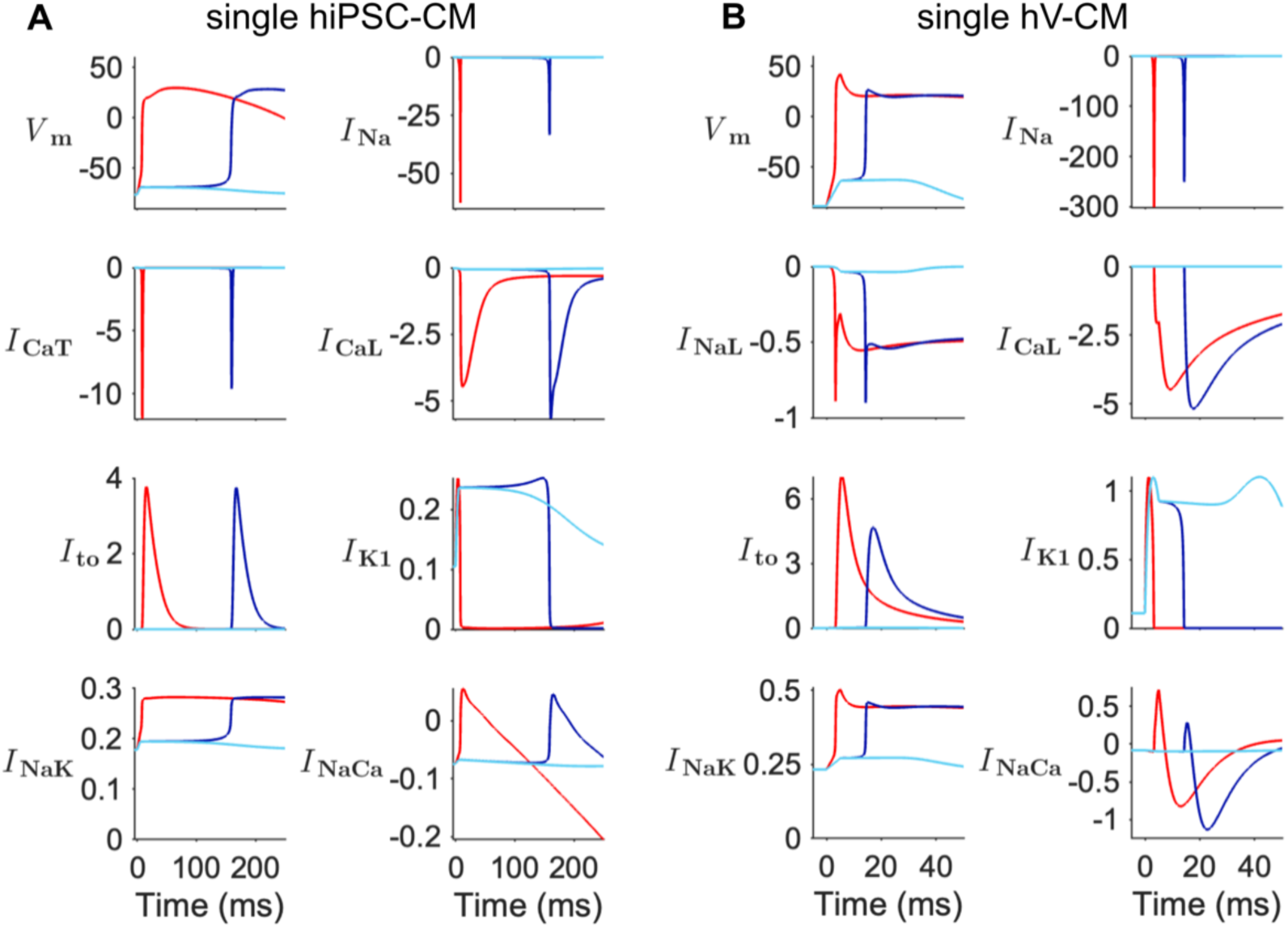
Single cell ionic currents during excitation with different stimulus strengths. (**A**) hiPSC-CM (mKC1) model. (**B**) hV-CM (ToR-ORd) model. Stimulus strength was either below threshold (cyan trace), at threshold (blue trace), or twice threshold (red trace). See **Fig. 3** legend for stimulus conditions. Note the different scales for the ordinates and time scales for hiPSC-CM vs. hV-CM. *V*_m_ has units of mV; currents have been normalized to cell capacitance with units of A/F.

For both hiPSC-CMs and hV-CMs, the largest currents active during the AP upstroke are *I*_Na_, *I*_CaL_, and *I*_to_ (and *I*_CaT_ for hiPSC-CMs). For both cell types, the magnitude of peak *I*_Na_ at threshold is smaller than at twice threshold, whereas the magnitude of peak *I*_CaL_ is actually larger. Also, peak *I*_CaT_ is smaller in hiPSC-CMs at threshold than at twice threshold. The main currents flowing during the sub-threshold response are *I*_K1_ and *I*_NaK_ in both cell types. Note that *I*_NaCa_ and *I*_CaL_ in hV-CMs (ToR-ORd model) consist of myoplasmic and subspace components, which have been combined here for comparison with *I*_NaCa_ and *I*_CaL_ in hiPSC-CMs. Furthermore, *I*_CaL_ in hV-CMs and hiPSC-CMs contains multiple ionic components dominated by Ca^2+^, which have been combined for comparison.

#### 2.2. Stimulus threshold for the strand

We define the stimulus threshold for the strand to be the minimum stimulus strength (*I*_T_) required to produce a propagating AP when the first five cells of the strand are excited (each with equal current) intracellularly with a 5 ms rectangular current pulse. Strands were first conditioned with a 1 Hz pulse train for 100 s, like for the single cell. Conduction velocity (*θ*) was constant (to within 1%) from 25% to 75% of the length of the strand, where boundary effects from the ends were absent, so it was measured during the 101^st^ beat at cell 35. *R*_i_ was set to 4 MΩ for the hiPSC-CM (mKC1) strand so that *θ* was 8.2 cm/s at 1000 ms pacing CL, which falls within the range of our experimental values from 2.7 to 18.1 cm/s (**Table 1**). *R*_i_ was also set to 1.67 MΩ for the hV-CM strand, so that *θ* was 48.4 cm/s, which falls within the range of reported values of 30 to 65 cm/s in transmural wedges and a 3D model of the human ventricle (Glukhov *et al*., 2010; Hyde *et al*., 2015). Values of *θ* at the other three pacing rates are given in **Appendix Table A1**, and those for the hiPSC-CM strand fall within our experimentally observed ranges at those rates (**Table 1**).

*I*_T_ was measured for the 101^st^ beat and is defined to be the minimum stimulus strength required to elicit a propagating AP with a peak voltage ≥ 0 mV at cell 50. *I*_T_ of the hiPSC-CM strand (**Fig. 5A**) is significantly larger than that of the single cell (4.20 pA/pF vs. 1.49 pA/pF). At threshold, an AP is elicited and propagates down the length of the strand. The initial activation delay following the break of the stimulus dissipates by around cell 8 for the hiPSC-CM strand and around cell 5 for the hV-CM strand, after which the activation time of the cell is purely from propagation delay. Additionally, even for a subthreshold stimulus, a small wave can spread down the entire length of the strand although its amplitude decays with distance (decremental conduction, red arrow, **Fig. 5A**). The fact that it can persist over such a long distance can be attributed to the elevation of *V*_m_ during diastole (due to intrinsic automaticity), approaching but not exceeding *V*_T_ of the cells and then, sustaining a graded response. However, if the maximal conductance of *I*_K1_ (*g*_K1_) is increased (e.g., by a factor of 5), the diastolic potential of the cell becomes more negative, its intrinsic automaticity is inhibited, and the subthreshold wave dies out by cell 30 (not shown). With stimuli above threshold, APs propagate down the strand, but with a shortened *t*_lag_, which for cell 1 of the strand is ∼56 ms at threshold and ∼4 ms at twice threshold. These times are less than the corresponding *t*_lag_s in the single cell. Thus, although the strand is more difficult to excite than the single cell, once excited, cells within the strand are excited more robustly.

**Figure 5.**
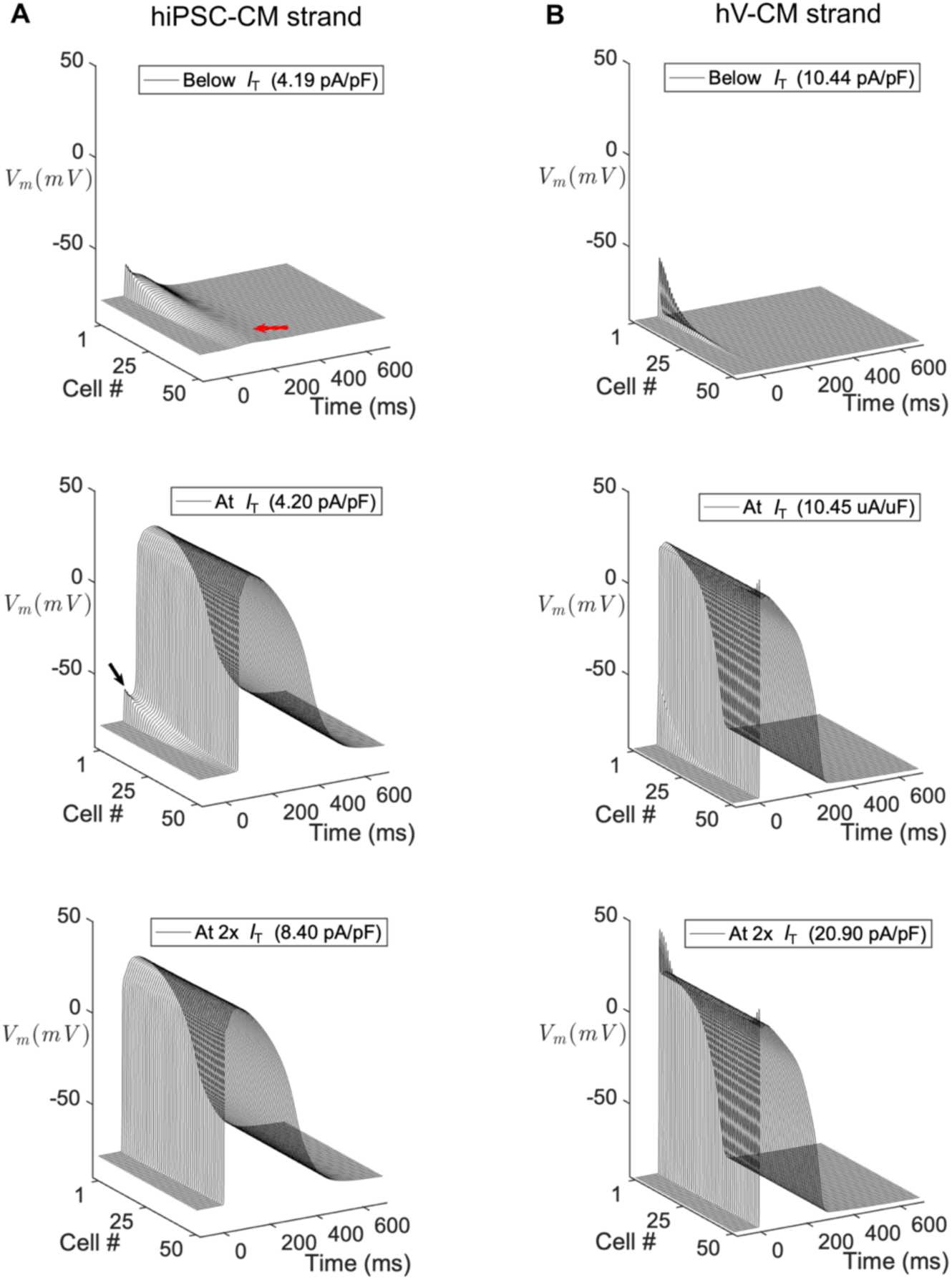
Strand excitation at different stimulus strengths. The strand was conditioned by a stimulus train of 100 pulses (1000 ms CL), after which it was stimulated 1 s later at its initial end either just below threshold (top row), at threshold, (middle row) or at twice threshold (bottom row). Shown are the *Vm* responses along the strand. (**A**) hiPSC-CM (mKC1) strand (red arrow indicates the decaying spread of the subthreshold depolarizing wave along the strand; black arrow indicates the threshold potential and lag time until the AP upstroke); (**B**) hV-CM (ToR-ORd) strand.

The hV-CM (ToR-ORd) strand (**Fig. 5B**) exhibits somewhat different behavior. Like for the hiPSC-CM strand, *I*_T_ of the ToR-ORd strand is larger than that of the single cell (8.81 μA/μF vs 5.81 μA/μF). However, at threshold the stimulated AP propagates without a large *t*_lag_. Below threshold, an active response is absent, and the subthreshold wave dies out by cell 40. *t*_lag_ in cell 1 of the strand is ∼8 ms at threshold and ∼2 ms at twice threshold, again less than *t*_lag_ in the single cell. Altogether, these results show that like hiPSC-CM strands, hV-CM strands are more difficult to excite than single hV-CMs but once excited, have more robust excitation. Furthermore, compared with hiPSC-CM strands, hV-CM strands have weaker decremental conduction and more robust propagation.

Shown in **Fig. 6** are the ionic currents in cell 1 for stimuli having subthreshold, threshold and twice threshold strengths.

**Figure 6.**
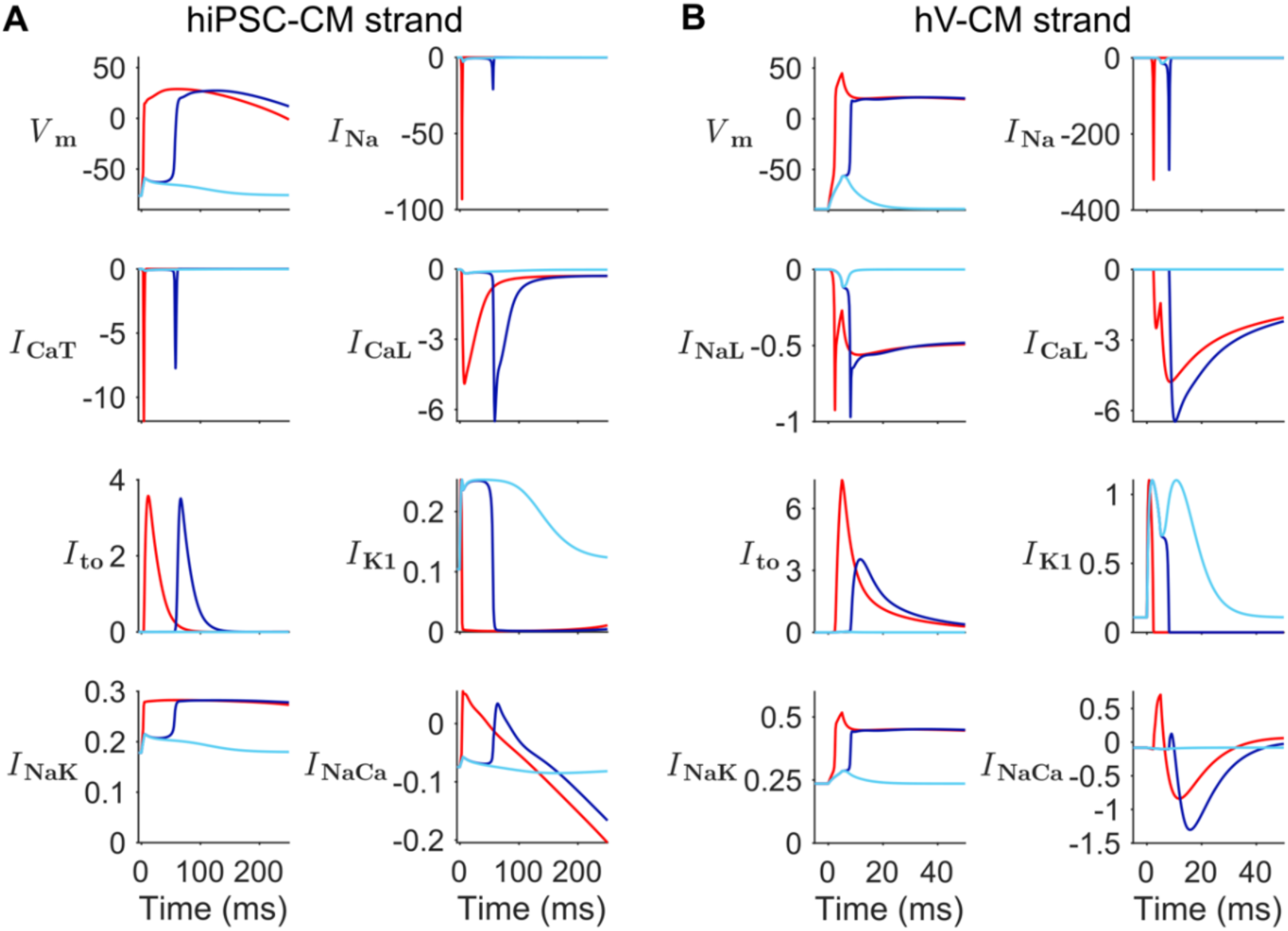
Strand excitation at different stimulus strengths during the depolarization phase. APs. (mV) and underlying ionic currents of cell 1. (**A**) hiPSC-CM (mKC1 model). (**B**) hV-CM (ToR-ORd model). Stimulus strength was either below threshold (cyan trace), at threshold (blue trace), or twice threshold (red trace). See **Fig. 3** legend for stimulus conditions. The largest ionic currents (*I*Na, *I*CaL, *I*to, *I*K1, *I*NaK, and *I*NaCa) are plotted below the APs. Note the different scales for the ordinates and time scales for hiPSC-CM vs. hV-CM. Units of *V*_m_ is mV and current is A/F.

We see that the patterns of currents are similar to those of the single cell for both cell types (**Fig. 4**), but with some distinct differences. Like for the single cell, the largest currents active during the AP upstroke are *I*_Na_, *I*_CaL_, and *I*_to_ for both hiPSC-CMs and hV-CMs, and additionally, *I*_CaT_ for hiPSC-CMs. For both cell types, peak *I*_Na_ at threshold is smaller but peak *I*_CaL_ is larger than at twice threshold, and for hiPSC-CMs, peak *I*_CaT_ is also larger. The main currents flowing during the subthreshold response are *I*_K1_ and *I*_NaK_. When stimulating at threshold, *t*_lag_ is much shorter and *V*_T_ is more positive than in the single cell. Also, at threshold *I*_Na_ is approximately 38% smaller in the hiPSC-CM strand than that of the single cell but is slightly larger (∼17%) for hV-CM strand vs. cell. Although *I*_CaL_ is small compared with *I*_Na_, it serves as a depolarization reserve in both cell types along with *I*_CaT_ in hiPSC-CMs that enables slow propagation to occur in situations in which *I*_Na_ is compromised. Note that the differences in ionic currents for threshold vs twice threshold stimulation are described here for cell 1 of the strand, and that these differences dissipate by the time propagation reaches the middle of the strand.

#### 2.3. Variation of stimulus threshold with stimulation rate

The simulations of **Figs. 3** and **5** were expanded to include pacing at 500, 700 and 2000 ms cycle lengths, and the values of *I*_T_, *V*_T_, and *V*_m0_ (*V*_m_ at the onset time of the stimulus pulse) with a 5 ms pulse were tabulated for the single hiPSC-CM, single hV-CM and their respective strands (**Table 3**).

**Table 3.** Stimulus threshold parameters for single cell and strand. Listed are values for *I*T, *V*T, *V*m0, and (d*V*m/dt)max for hiPSC-CM and hV-CM single cells and strands at four different pacing CLs (500, 700, 1000, 2000 ms). Parameters for the strands were measured at cell 1. Values for *I*T and (d*V*m/dt)max changed slightly in the second decimal place for hV-CMs if simulations were run beyond 100 beats.

| | $I_T$ (A/F) | | | | $V_T$ (mV) | | | | $V_{m0}$ (mV) | | | | $dV_m/dt$ (max) (V/s) | | | |
| --- | --- | --- | --- | --- | --- | --- | --- | --- | --- | --- | --- | --- | --- | --- | --- | --- |
| Pacing CL | 500 | 700 | 1000 | 2000 | 500 | 700 | 1000 | 2000 | 500 | 700 | 1000 | 2000 | 500 | 700 | 1000 | 2000 |
| single hiPSC-CM | 1.94 | 1.85 | 1.49 | 0.71 | -66.0 | -67.5 | -69.1 | -71.2 | -75.4 | -76.5 | -76.4 | -74.7 | 12 | 22 | 36 | 50 |
| hiPSC-CM strand | 5.14 | 4.79 | 4.20 | 2.93 | -58.1 | -61.2 | -63.1 | -68.7 | -75.2 | -76.5 | -76.4 | -74.7 | 6 | 9 | 20 | 36 |
| single hV-CM | 5.93 | 5.87 | 5.83 | 5.72 | -63.4 | -63.5 | -63.6 | -63.8 | -88.8 | -88.9 | -88.9 | -88.7 | 229 | 239 | 251 | 223 |
| hV-CM strand | 10.57 | 10.51 | 10.45 | 10.30 | -56.6 | -56.8 | -56.9 | -57.0 | -88.9 | -89.0 | -88.9 | -88.7 | 184 | 189 | 194 | 204 |

*I*_T_ is larger for the strand than for the single cell, as seen previously in **Fig. 5**, and this can be attributed to the loading effect of the electrically coupled cells downstream of the stimulated cells and subsequent reduction in excitability. *I*_T_ also increases for both the single cell and strand as pacing CL decreases (higher pacing rate), but by a greater amount (as a percentage change) for hiPSC-CMs than for hV-CMs, largely because the hiPSC-CM AP has not fully repolarized due to its long repolarization tail, resulting in residual refractoriness. *V*_m0_ is essentially the same in the strand as it is in the single cell, but *V*_T_ is more positive, again reflecting a lower excitability of the strand. Note that *V*_T_ becomes more negative along the strand from cell 1 to 50 (**Fig. 5A**), indicating increasing excitability. Also, *I*_T_ of hiPSC-CMs is smaller than that of hV-CMs for both single cells and strands at all four pacing rates, and *V*_T_ for hiPSC-CMs is generally more negative than that of hV-CMs. These differences reflect a greater excitability of hiPSC-CMs compared with hV-CMs.

#### 2.4. Variation of stimulus threshold with intracellular axial resistance

Intracellular axial resistance (*R*_i_), which is comprised of myoplasmic resistance (*R*_myo_) and gap junctional resistance (*R*_gap_), can change if gap junctional resistance were to increase, for example, under disease conditions. The pacing protocol of the previous section was applied, but with *R*_i_ set to different values during both the conditioning interval of stimulation and the test pulse. As *R*_i_ increases, both *I*_T_ and *θ* decrease monotonically (**Fig. 7**), consistent with previous studies (Shaw & Rudy, 1997).

**Figure 7.**
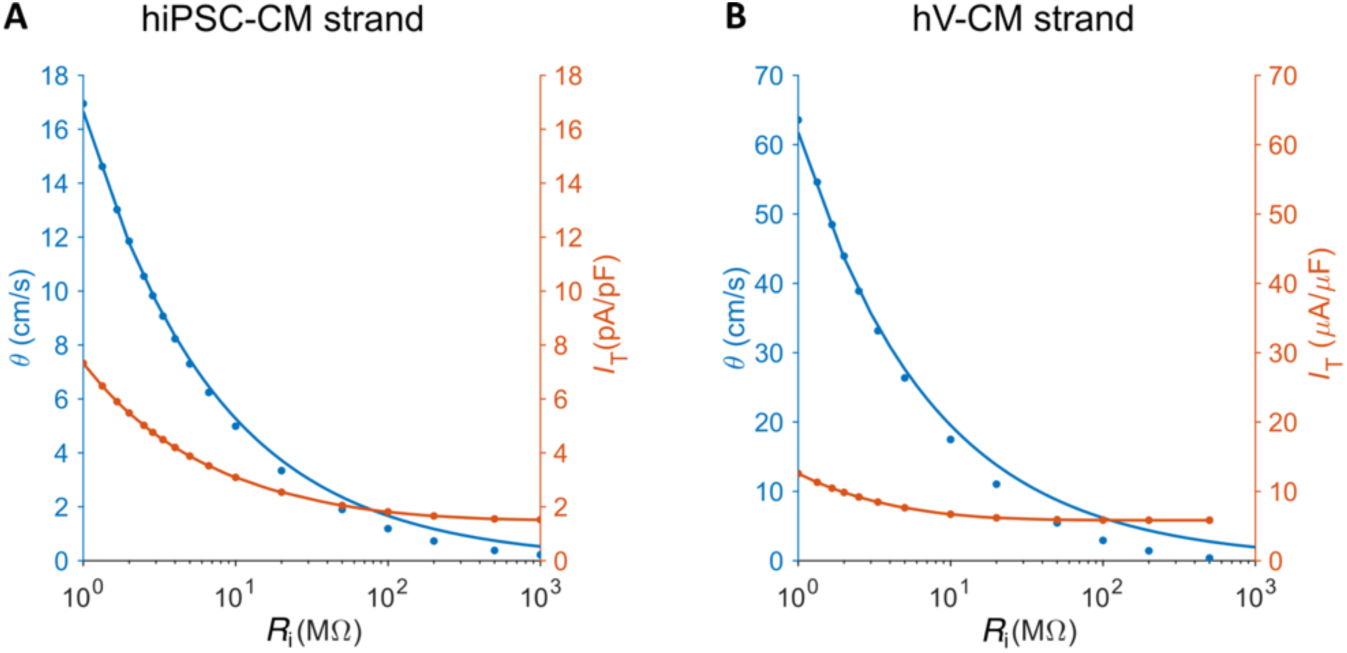
Dependence of stimulus threshold and conduction velocity on intracellular resistance. *I*T (red) and *θ* (blue) vs *R*_i_ (note log scale) for (**A**) hiPSC-CM strand (mKC1 cable) and (**B**) hV-CM strand (ToR-ORd cable). The solid blue trace was fit according to **Eq. 3** with the constant *k* having a value of 1.5 x 10^6^ MΩ-cm-s^-2^ for the hiPSC-CM strand and 7.7 x 10^6^ MΩ-cm-s^-2^ for the hV-CM strand.

At very high R_i_ (∼1000 MΩ), e.g., with very low gap junctional coupling, *θ* approaches 0, but *I*_T_ approaches a non-zero value, as expected theoretically (Geddes & Bourland, 1985). It is well known that in a one-dimensional cable with high extracellular conductivity, *θ* varies inversely with the square root of *R*_i_ (Hodgkin, 1954).

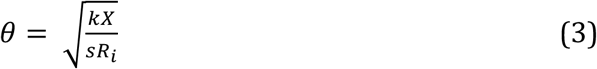

where *X* is the cross-sectional area, *s* is the membrane area per unit length, and *k* is a constant that depends on the local properties of the membrane, such as species, type of membrane and temperature (Plonsey & Barr, 2007). The conduction data in **Fig. 7** are fit quite well by **Eq. 3** for *R*_i_ up to around 10 MΩ, above which significant divergence occurs because the cells become sufficiently decoupled such that their behavior deviates from that of a continuous cable.

### 3. Strength-duration relations for the single cell and strand

#### 3.1. Strength-duration curves

Electrical stimulation of excitable cells and tissue by a current source follows a well-known inverse relation between the strength of the stimulus pulse at stimulus threshold and its duration. This was characterized mathematically by Weiss (Weiss, 1901) and later by Lapicque (Lapicque, 1907) who defined two parameters – the *rheobase* (the minimal stimulus strength *I*_rh_ below which excitation cannot occur no matter how long the stimulus duration *d*), and *c*, the *chronaxie* (the stimulus duration having a threshold strength equal to twice the rheobase). For a rectangular stimulus pulse, the threshold current (*I*_T_) and the threshold charge (*Q*_T_, equal to the time integral of *I*_T_) are:

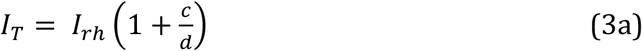

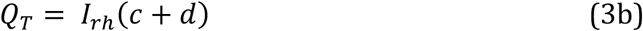

A variant of these relations is based on a passive RC membrane charging to a threshold voltage, and takes the exponential forms:

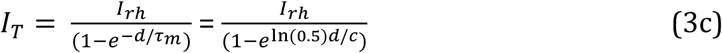

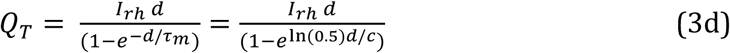

where *τ*_m_ is the membrane time constant equal to the product of effective membrane resistance *R*_m_ and capacitance *C*_m_ and can been replaced by *c/*ln(0.5).

For simplicity, we will focus on **Eqs. 3a** and **3b**. Note that **Eqs. 3** pertain to electrical stimulation by a transmembrane current source, and that external stimulation of a cell by electrical fields involves a different biophysical process in which different segments of cell membrane are simultaneously hyperpolarized and depolarized (Tung & Borderies, 1992). The strength-duration relationship has been used to systematically evaluate factors influencing the electrical excitation of animal ventricular cells in vitro such as cell orientation and dimensionality of tissue (Tung *et al*., 1991; Sayegh *et al*., 2019; Li *et al*., 2020).

The stimulus threshold presented in the previous section for *d* of 5 ms at a pacing CL of 1000 ms can be determined for other pulse durations to generate the strength-duration relations for current and charge (**Fig. 8**). The data points in **Fig. 8B** fall on nearly a straight line and have a better (lower root mean square error, RMSE) fit with the linear relation (**Eq. 3b**) than with the exponential relation (**Eq. 3d**), particularly for the hV-CM strand.

**Figure 8.**
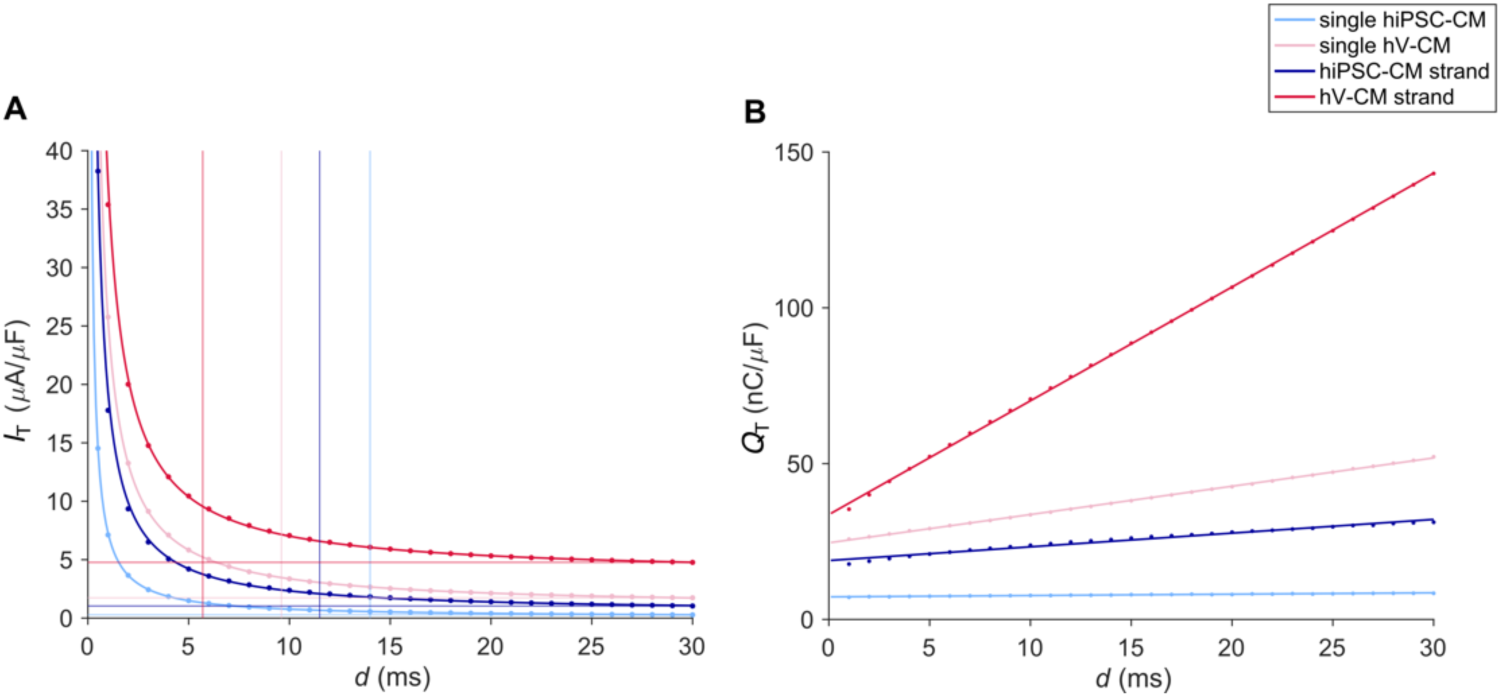
Strength-duration relations for the hiPSC-CM and hV-CM single cell and strand. For the strands, the same stimulus current was applied to cells 1-5. **A.** Stimulus current. Horizontal lines indicate values of rheobase, while vertical lines indicate values of chronaxie estimated at *d* = 30ms. Solid curves are based on **Eq. 3a** using parameters from fits in (**B**). All currents have been normalized to cell capacitance. **B.** Stimulus charge. Straight lines are best fit of **Eq. 3b** to the data points.

For both cell types, the threshold currents for the strands are generally greater than those for the single cell, because of the loading effect of downstream cells. If the number of cells being stimulated is increased, this counters the loading effect, and the SD relation for the strand shifts downward, asymptotically approaching that of the single cell (**Appendix Fig. A3**). Also, with *g*_Na_ block or higher pacing rates, the SD relations all shift upward (not shown), with the latter reflecting a process of rate-accommodation due to inactivation of *I*_Na_ (Davidenko *et al*., 1990). Lastly, the SD relations for the hiPSC-CM cell and strand are lower than the corresponding relations for the hV-CM cell, which reflects the greater excitability of hiPSC-CMs. The relative differences among the SD relations for current and their shifts with load, *g*_Na_ block, or pacing rate are similar for the SD relations for charge.

#### 3.2. Relationship of rheobase and chronaxie to individual ionic currents

Next, we examined whether the characteristic parameters of the SD relation, namely chronaxie and rheobase, depend strongly on individual ionic currents. The curves in **Fig. 8B** were fit with **Eq. 3b**, yielding values for *I*_rh_ and *c* (**Table 4A**).

**Table 4.**
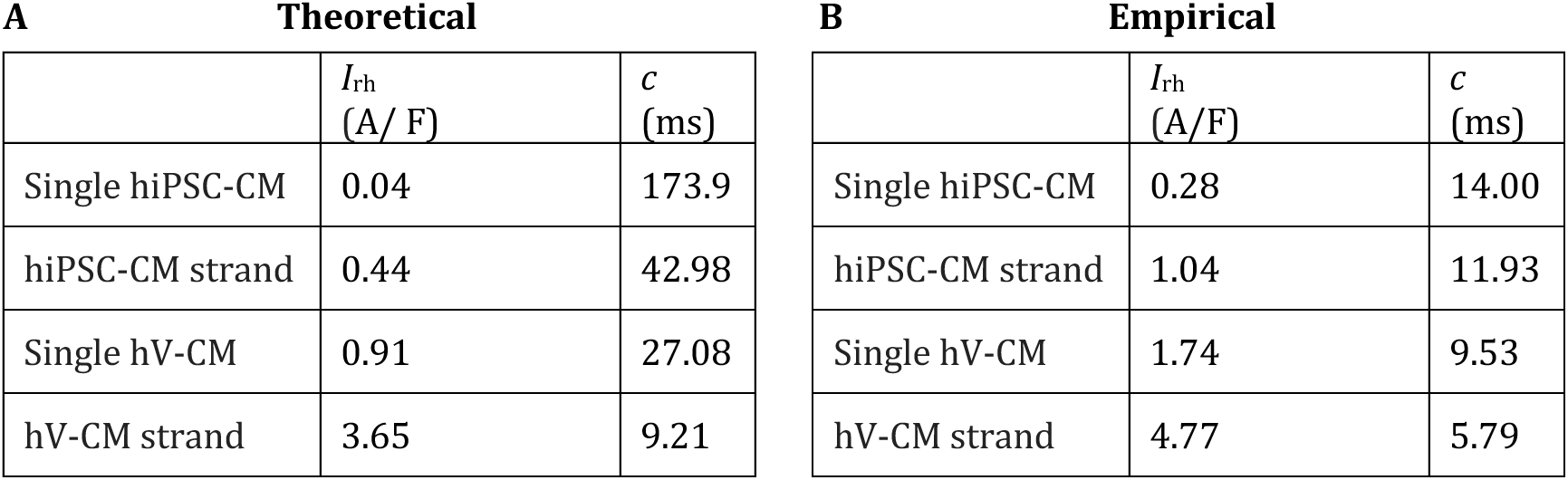
Theoretical rheobase and chronaxie estimated from **Eq. 3b**. **B.** Rheobase and chronaxie values estimated empirically at *d*= 30ms for the strength-duration relations shown in **Fig. 8A**. All currents have been normalized to cell capacitance.

| A Theoretical |  |  | B Empirical |  |  |
| --- | --- | --- | --- | --- | --- |
| | $I_{rh}$<br>(A/ F) | $c$<br>(ms) | | $I_{rh}$<br>(A/F) | $c$<br>(ms) |
| Single hiPSC-CM | 0.04 | 173.9 | Single hiPSC-CM | 0.28 | 14.00 |
| hiPSC-CM strand | 0.44 | 42.98 | hiPSC-CM strand | 1.04 | 11.93 |
| Single hV-CM | 0.91 | 27.08 | Single hV-CM | 1.74 | 9.53 |
| hV-CM strand | 3.65 | 9.21 | hV-CM strand | 4.77 | 5.79 |

Because it is experimentally difficult to stimulate with a very long pulse (but possible with the computational models), the rheobase is sometimes defined empirically as *I*_T_ at some pulse duration. For example, one study used *d* = 2 ms (Kay *et al*., 1990) but here, we will use *d* = 30 ms (as shown with vertical lines in **Fig. 8A**), which then establishes the values for *c* (horizontal lines in **Fig. 8A**). These values for *I*_rh_ and *c* are given in **Table 4B** and differ substantially from the theoretical values, suggesting that empirical measurements of rheobase may be fraught with error.

We then performed a sensitivity analysis of the theoretical *I*_rh_ and *c* as defined in **Eqs. 3**, as well as *I*_T_ for a 5 ms pulse, at multiple pacing rates in both hiPSC-CMs and hV-CMs to both a 10% increase and a 10% decrease in conductance of a given ionic current (**Fig. 9**). The only exception was for *I*_Kr_ in hiPSC-CMs, where a 10% increase resulted in repolarization failure, so *I*_Kr_ was changed by only 5%.

**Figure 9.**
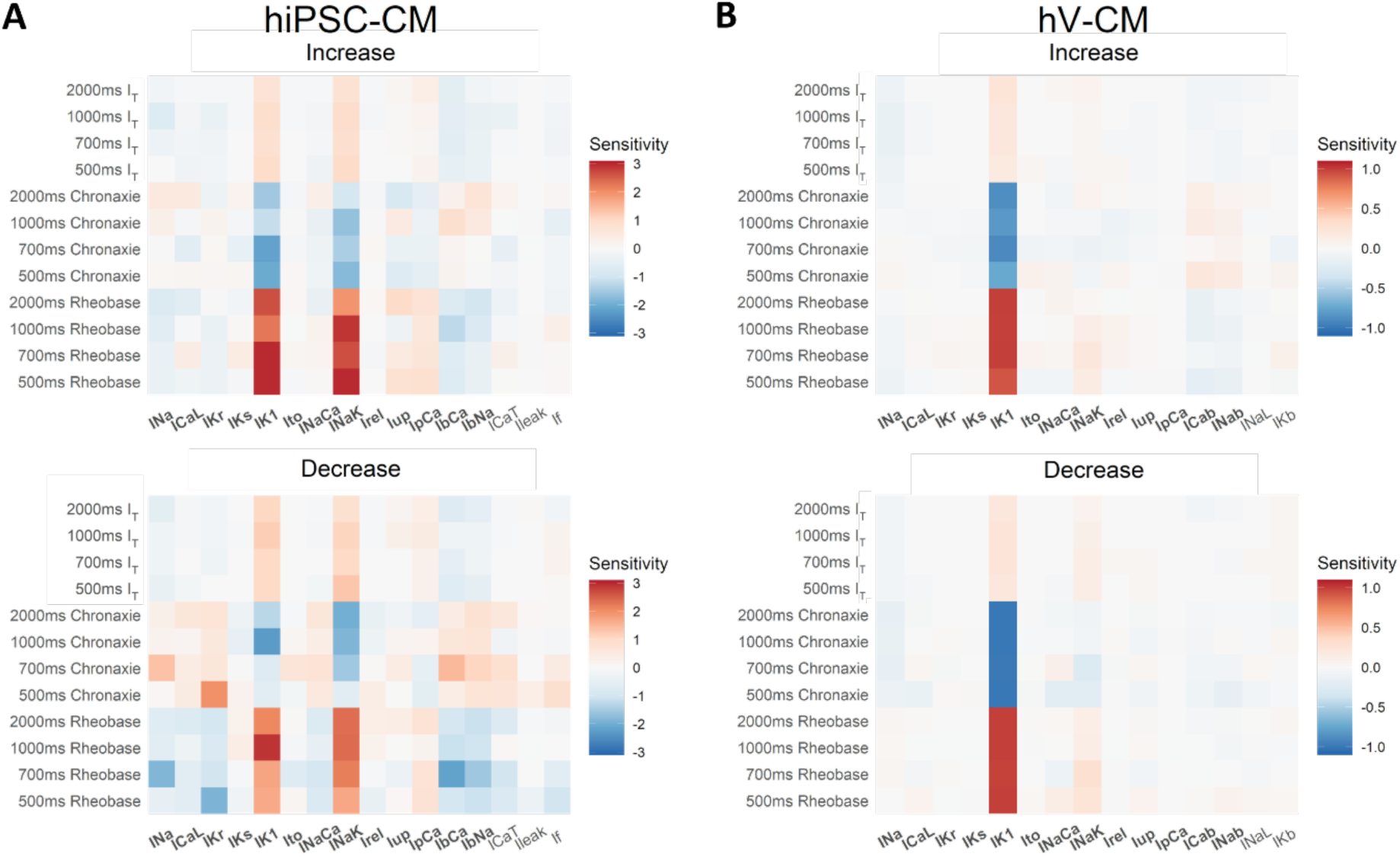
Sensitivity analysis of single hiPSC-CM (mKC1 model) and single hV-CM (ToR-ORd model). (Upper row) Heatmaps showing changes in stimulus threshold (for a 5 ms pulse), rheobase and chronaxie at four pacing cycle lengths when conductance values for ionic currents were individually perturbed by +10% (with the exception of 5% for *IKr* in the mKC1 model). (Lower row) Heatmaps for *I*T, rheobase and chronaxie when conductance values were individually perturbed by –10%. Values exceeding the displayed ranges were assigned the color at the ends of the ranges. Currents shared between the two cell types are boldfaced (all but the last two or three on the right). Sensitivity values in the heatmaps are the ratio of change in the output (*I*T, *c*, or *I*rh) to the change in the given conductance. Positive values are shown in red, and negative values in blue.

The sensitivity pattern is rather clear for hV-CMs: *I*_T_, *c*, and *I*_rh_ are strongly and mainly dependent on *I*_K1_, with sensitivities that are relatively unaffected by the polarity of the conductance change. However, in hiPSC-CMs, while *I*_T_, *c*, and *I*_rh_ also depend strongly on *I*_K1_, they depend on *I*_NaK_ (sodium-potassium pump) as well, with sensitivities that are generally greater for an increase rather than for a decrease in the conductances. *c* and *I*_rh_ are also moderately sensitive to a decrease in conductance of several of the other ionic currents, including *I*_Na_, *I*_Kr_, *I*_bCa_, and *I*_bNa_.

Sensitivity maps of the *empirical* values of *I*_T_, *c* and *I*_rh_ (not shown) are qualitatively similar to those in **Fig. 9** but tend to be lower in amplitude for *c* and *I*_rh_. In addition, for hiPSC-CMs, *c* becomes insensitive to these two currents. For hV-CMs, *c* becomes slightly less sensitive to *I*_K1_ and more sensitive to *I*_NaK_.

#### 3.3. Variation of excitatory currents with different stimulus pulse durations

Lastly, we examined the process of excitation in the single cell for the various ionic currents at different stimulus pulse durations – specifically, for stimuli at twice threshold for a duration *d* of 1, 5 or 20 ms. The results are shown in **Fig. 10**.

**Figure 10.**
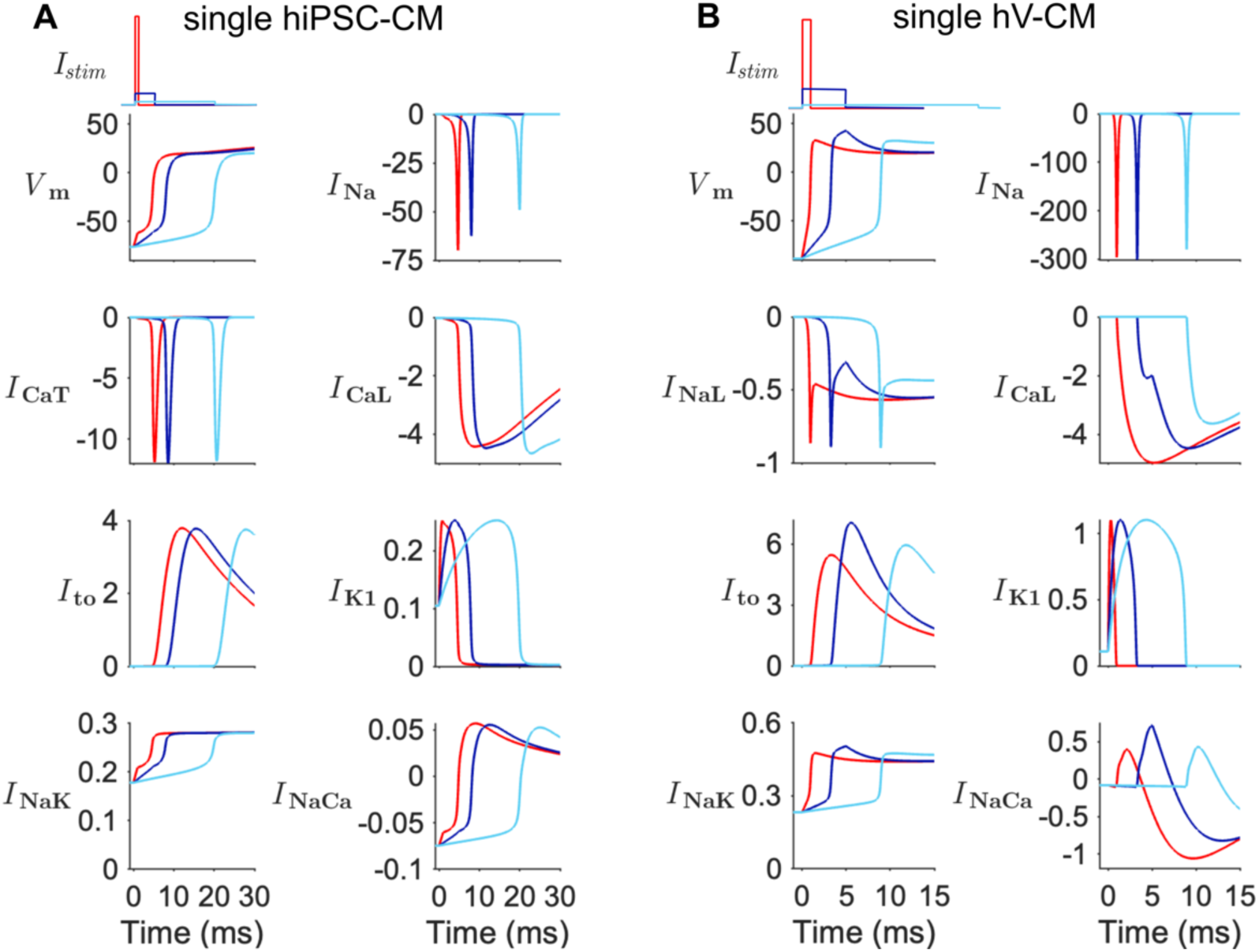
Single cell excitation with different stimulus pulse durations. APs. (mV) and underlying ionic currents are shown for **A.** hiPSC-CM (mKC1 model) and **B.** hV-CM (ToR-ORd model). See **Fig. 3** legend for stimulus conditions. The APs and their largest ionic currents (*I*Na, *I*CaL, *I*to, *I*K1, *I*NaK, and *I*NaCa) are plotted.

Stimulus amplitude at each pulse duration (1 ms in red, 5 ms in blue, and 20 ms in cyan) was set at twice threshold for that pulse duration. Note the different scales for the ordinates. Units of *V*_m_ is mV and current is A/F.

We see that as *d* increases from 1 to 20 ms, *t*_lag_ increases substantially, and the peak amplitude of *I*_Na_ decreases by ∼30% in hiPSC-CMs and ∼6% in hV-CMs. This decrease can be attributed to the increased time for *V*_m_ to reach *V*_T_, which increases the degree of inactivation (from gating variable *h* but not *j*) of *I*_Na_. Despite these reductions, there is still sufficient *I*_Na_ to excite the cell. Increased *d* also increases *I*_CaL_ by ∼5% in hiPSC-CMs but decreases *I*_CaL_ (both subspace and cytosolic components) by ∼27-28% in hV-CMs. The amplitudes of the other currents remain relatively unchanged with increasing *d*, except for *I*_to_, *I*_NaK,_ and *I*_NaCa_ in hV-CMs, which increase and then decrease in amplitude.

### 4 Cell within the strand vs single cell

#### 4.1. Stimulation by propagating action potential

Cells within a strand are excited not by an exogenous current source but by a propagating AP. In any finite length strand, the cells at the two ends tend to have boundary effects (evident in the larger overshoots of their APs in **Fig. 5B**) and are not representative of cells in the middle. Among the 50 cells in our strands, cells 30-40 were found to have similar activation (phase 0) time courses; hence cell 35 was chosen to be the representative cell within the strand. AP shape and all of the ionic currents underlying cell 35’s excitation are essentially unchanged (apart from a shift in activation time) regardless of whether the strand’s end is stimulated at threshold, at twice threshold, or with different pulse durations, although slight variations in *θ* were observed. The ionic currents for cell 35 are shown in **Fig. 11A** for the hiPSC-CM strand and in **Fig. 11B** for the hV-CM strand.

**Figure 11.**
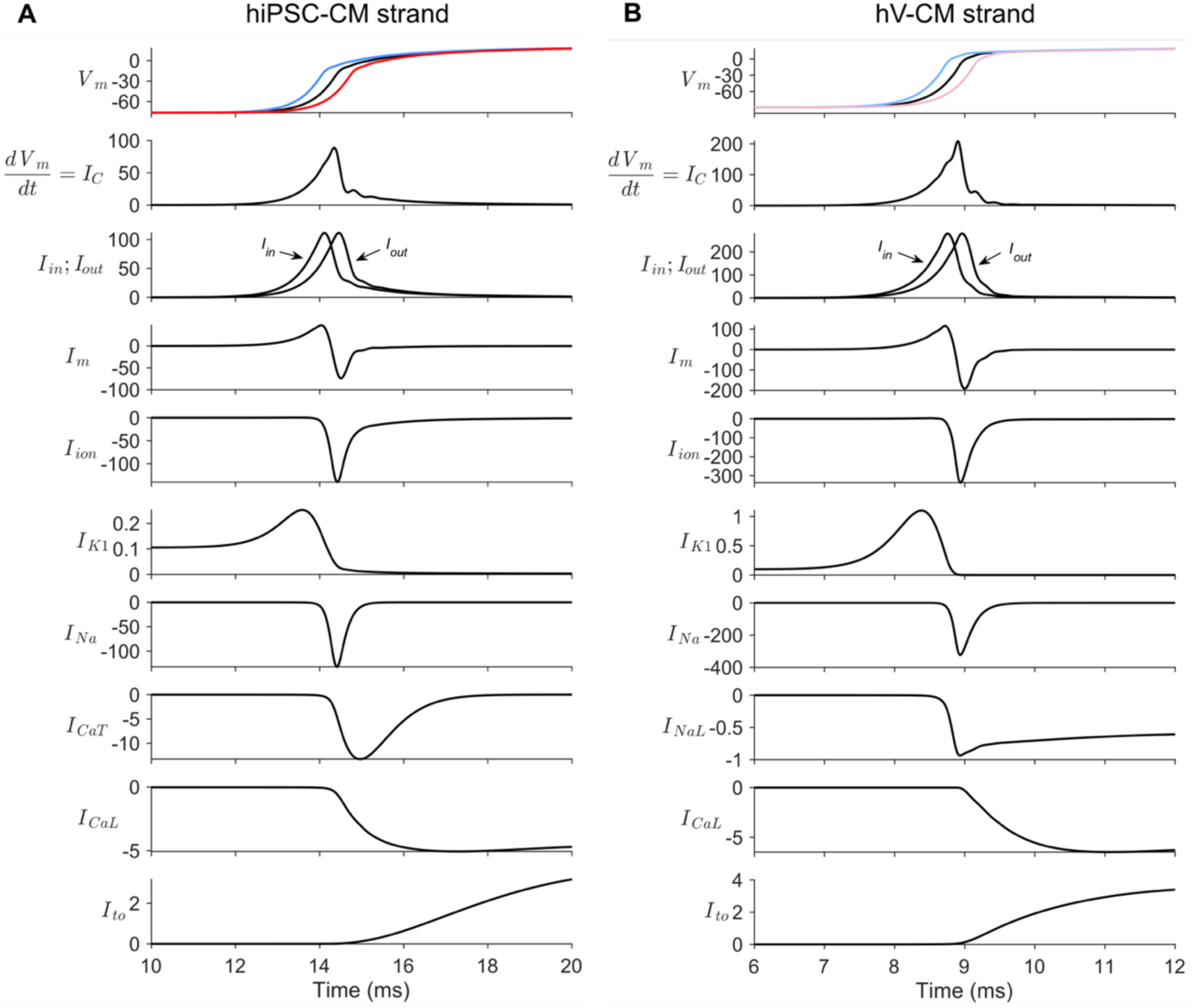
Excitation of cells 34-36 in a strand by a propagating AP. **A.** hiPSC-CM strand (mKC1 cable) with *R*_i_ = 4 MΩ. **B.** hV-CM strand (ToR-ORd cable) with *R*_i_ = 1.67 MΩ. Top row: APs (mV); second row: *dV*_m_/*dt* (equal to *I*c in **Eq. 2**; mV/ms); remaining rows: ionic currents normalized to cell capacitance (A/F). Stimulus strength was 2x threshold applied to cable cells 1-5. Cell 34 (blue traces), cell 35 (black traces), cell 36 (red traces). Currents for cells 34 and 36 are not shown. Note the different scales on the ordinates.

As the AP progresses down the hiPSC-CM strand at a speed of 8.2 cm/s, there is a propagation delay of 0.34 ms between cells 34 and 35 (and between cells 35 and 36) and a (*dV*_m_/*dt*)_max_ of 89 V/s. In the hV-CM strand the wavefront moves more quickly at a speed of 48.4 cm/s with a shorter propagation delay per cell of 0.21 ms and a higher (*dV*_m_/*dt*)_max_ of 209 V/s. *I*_C_, *I*_in_, *I*_out_, *I*_m_, and *I*_ion_, as defined in **Fig. 1C** and in **Eqs. 1** and **2**, are shown in addition to the individual ionic currents. Compared with **Fig. 4**, the magnitudes of the ionic currents are similar to those for the single cell at twice threshold, with the exception of *I*_Na_, which is much larger than in the single cell. The reason for this increase, as discussed later, is from differences in the time course of the stimulating current. For the hV-CM strand, *I*_Na_ and *I*_CaL_ exhibit slightly higher magnitudes for the cell in the strand compared with the single cell, except for *I*_to_, which is slightly reduced.

When the cell lies in a strand, additional currents flow into and out of the cell with its neighbors. *I*_in_ is a monophasic current flowing into cell 35, and *I*_out_ is a monophasic current flowing out of the cell. A discussion of these currents can be found in (Shaw & Rudy, 1997; Gray *et al*., 2013) and is expanded upon here. *I*_m_ is the difference between *I*_in_ and *I*_out_ and is also equal to the sum of *I*_C_ and *I*_ion_ (**Eq. 1**). At cell 35, propagation has essentially reached steady-state, so *I*_out_ is essentially identical to *I*_in_ except for a time delay equal to the intercellular propagation delay. Being the difference between *I*_in_ and *I*_out_, *I*_m_ is necessarily biphasic. As the sum of membrane currents, *I*_m_ is mostly from *I*_C_ during the initial part of its positive phase and all from *I*_ion_ during its negative phase. *I*_ion_ is the sum of all the ionic membrane currents and is mostly *I*_K1_ until *V*_m_ reaches *V*_T_, mostly *I*_Na_ during the rapid *V*_m_ upstroke, and mostly either *I*_CaT_ (for hiPSC-CM) or inward *I*_NaL_ (for hV-CM) during the final phase of the upstroke and AP peak, with increasing contribution from *I*_CaL_.

#### 4.2. Stimulus current in the strand

The question now arises as to the current with which one can stimulate the single cell so that it responds as if it were in a strand. That current is none other than *I*_m_ (Shaw & Rudy, 1997; Gray *et al*., 2013), which is a biphasic current during the upstroke (**Fig. 11**) and has a net zero total area (charge) under steady-state conditions when integrated over an entire AP cycle. Note that the stimulus current is not *I*_in_, as some of *I*_in_ flows downstream to the next cell as part of *I*_out_. As a check, we stimulated the single hiPSC-CM and hV-CM with their corresponding *I*_m_ waveforms in **Fig. 11** and obtained the same AP and ionic currents shown in **Fig. 11** for cell 35.

For a 1D strand lying in a large conductive bath, *I*_m_ is closely proportional to the second spatial derivative of *V*_m_ along the strand, assuming continuous, uniform propagation (Plonsey & Barr, 2007), which in turn is proportional to the second temporal derivative of *V*_m_ according to the wave equation:

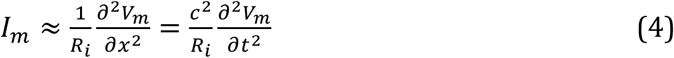

where *R*_i_ is the intracellular axial resistance per unit length (**Fig. 1**). As noted earlier, the assumption of a continuous cable, and hence **Eq. 4**, is not suitable at large values of *R*_i_.

During excitation, *I*_m_ can be approximated by an asymmetric biphasic rectangular or biphasic triangular pulse as shown in **Fig. 12A**. *V*_m_ and the ionic currents elicited by all three stimulus waveforms are compared in **Fig. 12B**.

**Figure 12.**
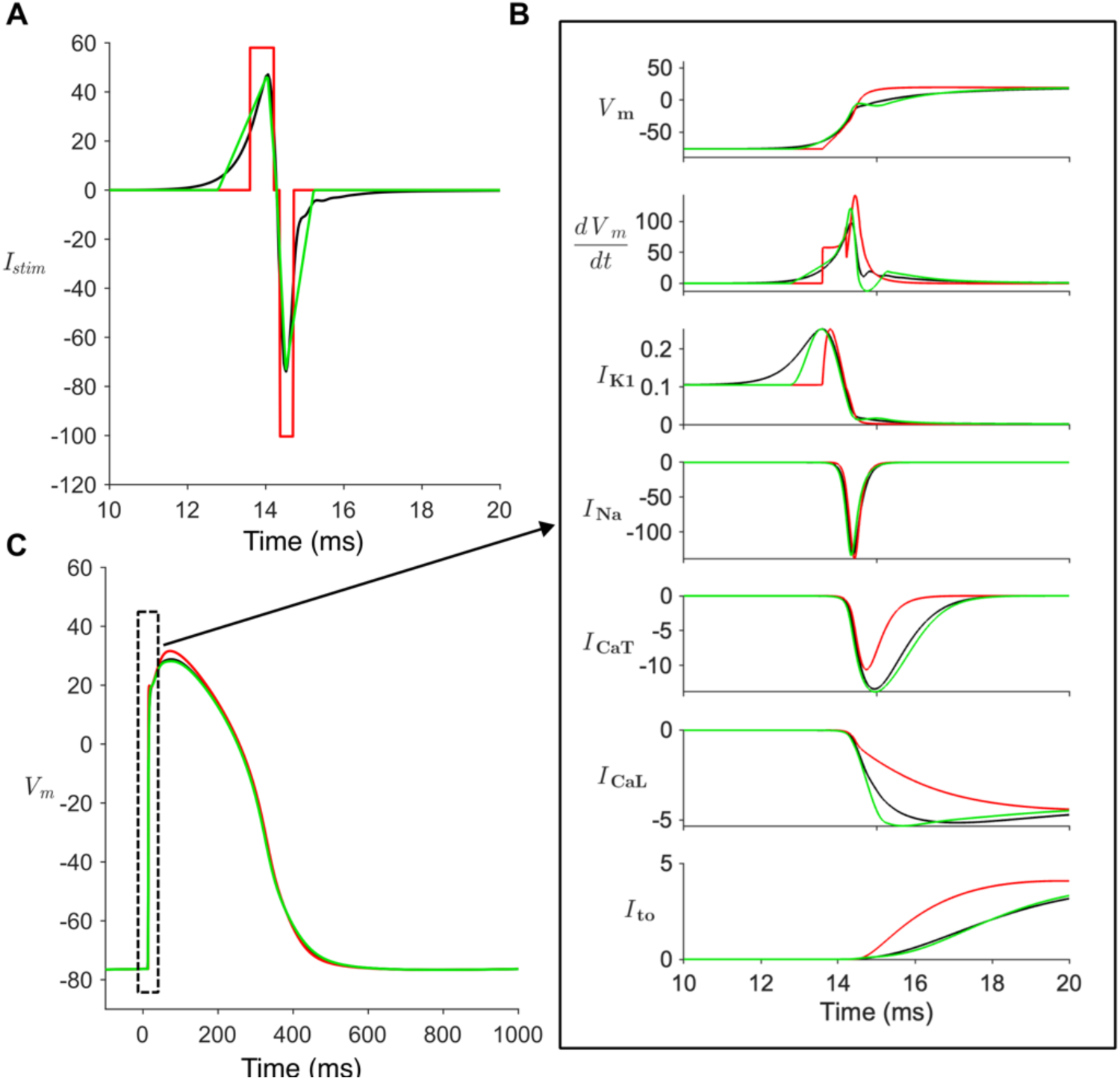
Excitation of single hiPSC-CM by three different stimulus waveforms. **A.** Stimulus waveforms are either *I*m for cell 35 in a strand (black), a biphasic rectangular pulse approximating *I*m (red), or a biphasic triangular pulse approximating *I*m (green). **B.** Voltages and ionic currents of the cell elicited by the three waveforms. **C.** APs over the entire pacing cycle, when elicited by the three waveforms. Units of *V*_m_ is mV, d*V*m/dt is mV/ms, and current is A/F.

From **Eq. 4**, the biphasic rectangular pulse implies that the upstroke of the AP is represented as two piecewise quadratic functions. We define the pulse to have an initial positive phase with a duration equal to the full width at half maximum (FWHM) of the positive half of *I*_m_’s initial spike. Its amplitude is such that its total charge, or area, equals that of the positive half of *I*_m_. Likewise, the latter negative phase has a duration equal to the FWHM of the negative half of *I*_m_’s initial spike and an amplitude such that its charge equals that of the negative phase of *I*_m_ up to its second zero crossing. It follows the end of the first phase by a short interphase interval. Stimulation with the biphasic rectangular pulse (**Fig. 12B**) yields nearly the same AP and *I*_Na_ as when stimulating with *I*_m_. However, *I*_CaT_ has smaller amplitude and faster decay, *I*_CaL_ activates more slowly, while *I*_to_ activates more rapidly. These differences in kinetics arise because of the step increase in current with the rectangular pulse, as opposed to the slower continuous change in amplitude with the physiological *I*_m_.

The biphasic triangular pulse allows the upstroke of the AP to be represented as two piecewise cubic functions, with continuity of slope between them. We define it to have an initial positive phase with an amplitude equal to the peak amplitude of the positive phase of *I*_m_, a duration such that its total charge equals that of the positive phase of *I*_m_ and terminates at the first zero crossing of *I*_m_. The latter negative phase begins at the zero crossing of *I*_m_, has an amplitude equal to the peak negative amplitude of *I*_m_, and a duration such that its charge matches that of the negative phase of *I*_m_ up to its second zero crossing. *V*_m,_ ionic currents and their gating variables are all very similar to those produced by *I*_m_, showing that the biphasic triangular waveform is a close and useful approximation of *I*_m_.

In summary, the physiological stimulation of cells by a propagating AP within a strand is by a complex biphasic pulse rather than by a monophasic rectangular stimulus pulse as used in the traditional strength-duration relations (**Fig. 8**). A biphasic triangular pulse is a good approximation to the physiological *I*_m_, although with diminishing fidelity for *R*_i_, > 10MΩ (not shown). On the other hand, a biphasic rectangular pulse is particularly erroneous, especially with respect to the kinetics of the activation of ionic currents which now undergo a step function rather than a monotonic increase. The challenge in generating a strength-duration curve for the biphasic triangular pulse is that three parameters (duration, peak amplitude, and timing of the peak) are required to characterize each phase of the waveform, giving a total of six independent parameters.

Nonetheless, we can address a separate question as to whether the latter negative phase of the biphasic physiological pulse has an inhibitory effect on the excitatory effect of the initial positive phase, given that it reduces the net charge contained in *I*_m_. **Fig. 13** compares stimulation by the initial, positive phase of *I*_m_ alone with that by the full biphasic *I*_m_.

**Figure 13.**
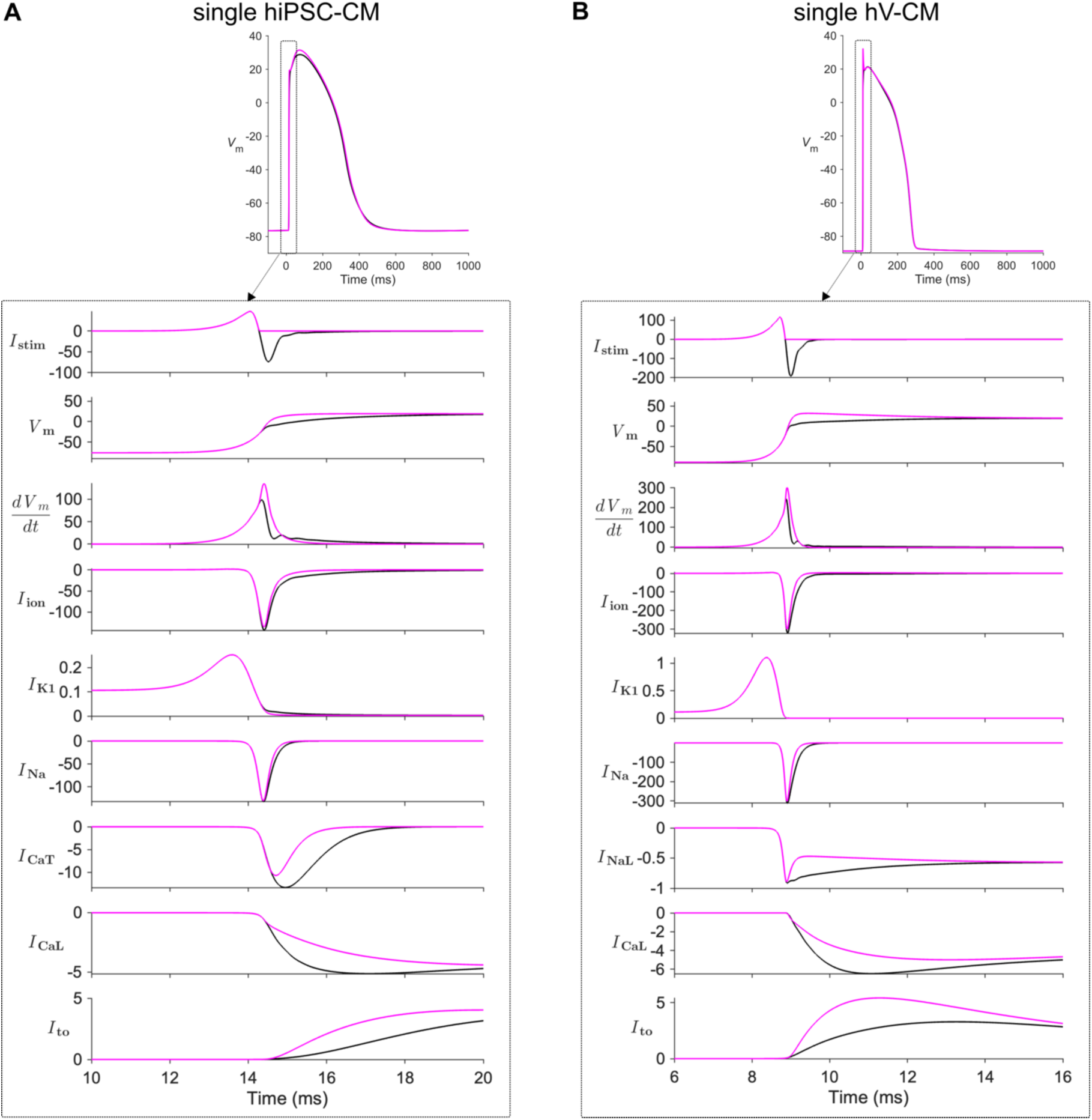
Excitation of single hiPSC-CM and hV-CM by monophasic or biphasic stimulus. Stimulus waveforms are either the initial, positive phase of *I*m for cell 35 in a strand (magenta) or the full biphasic *I*m (black), with *R*_i_ = 4 MΩ for the mKC1 cable or 1.67 MΩ for the ToR-ORd cable. **Upper row.** AP over the entire pacing cycle elicited by the two waveforms. **Lower rows.** Voltages and ionic currents of the cell elicited by the two waveforms. Portions of the black traces are hidden by the overlying magenta traces. Units of *V*_m_ is mV, d*V*m/dt is mV/ms, and current is A/F.

As expected, the latter negative phase of the biphasic pulse truncates the fast-rising phase of *V*_m_ with very little change in *I*_K1_. Perhaps non-intuitively, the subtle change in the upstroke phase results in a *larger* peak inward *I*_CaT_ (for hiPSC-CM) or *I*_NaL_ (for hV-CM), a *larger* peak inward *I*_CaL_ an incremental *increase* in peak *I*_Na_, and a *smaller* peak outward *I*_to_, all of which increase and prolong inward *I*_ion_. The time courses of the currents for monophasic and biphasic pulses converge after around *t* = 20 ms for both hiPSC-CMs and hV-CMs and thereafter are largely unaffected. Despite these changes in peak currents, the peak of the AP does not increase but actually decreases **(Fig. 13B)**, with little change in APD_30_ or APD_80_. This apparent paradox can be explained by recognizing that *I*_ion_ supplies current not only to charge the capacitance of cell 35, but also to charge the capacitances of downstream cells via *I*_out_. Thus, the latter negative phase of the physiological stimulus current augments the inward currents (*I*_Na_, *I*_CaT_. *I*_NaL_, and *I*_CaL_) and inhibits the outward currents (*I*_to_) activated by the initial positive phase via the alteration in *V*_m_ time course, thereby boosting the source current available to excite downstream cells.

The biophysical basis for the changes in these currents during the AP upstroke can be elucidated by examination of their gating variables. For hiPSC-CMs, the slight increase in peak *I*_Na_ is primarily from greater activation of the gating variable *m* that is not completely offset by greater inactivation of *h*. *I*_Na_ is also prolonged due to slower inactivation of *j* and an increase in electrochemical driving force for Na^+^. The increase in peak *I*_CaT_ is primarily due to greater activation of *dCaT*, delayed and slower kinetics of inactivation of *fCaT*, and an increase in the driving force for Ca^2+^. The increase in *I*_CaL_ arises mainly from an increase in the combined permeability and driving force for Ca^2+^of the calcium component of *I*_CaL_, *ibarca* that is only partially offset by slower activation of *d*. Finally, the decrease in *I*_to_ is primarily from slower and delayed activation of *r* and reduction in driving force for K^+^. For hV-CMs, the gating variables primarily responsible for the increases in peak *I*_Na_, *I*_NaL_, and *I*_CaL_ and decrease in peak *I*_to_ are respectively: greater activation of *m* despite greater inactivation of *h* and its phosphorylated form *hp*; greater activation of *mL*; slower inactivation of various slow, fast and phosphorylated components of *f* and of calcium-dependent *fCa,* together with an increase in the driving force for Ca^2+^; and slower activation of *a* and its phosphorylated form *ap*, greater inactivation of *i* and its phosphorylated form *ip*, and a smaller driving force for K^+^.

#### 4.3. Variation of *I*_m_ and ionic currents with pacing rate

As the pacing CL decreases in both strands, APD shortens, consistent with experimental observations. In addition, there is a significant increase in activation time (owing to slower *θ* and longer time for the AP to reach cell 35), a decrease in the magnitude and an elongation of *I*_m_, particularly in the hiPSC-CM strand but much less so in the hV-CM strand. *I*_ion_ also becomes smaller and elongated in the hiPSC-CM strand but is nearly unchanged in the hV-CM strand. The former is primarily a reflection of the temporal changes in *I*_Na_ and a reduction of its total charge by almost 50% when decreasing CL from 2000 to 500 ms. Most responsible for the *I*_Na_ changes in hiPSC-CMs is a rate-dependent decrease in *j* and secondarily, a decrease in *h* despite an increase in driving force for Na^+^. In hV-CMs, the rate-dependent decrease in *I*_Na_ is also from a decrease in the fraction of non-phosphorylated channels.

#### 4.4. Variation of stimulus and ionic currents with gap junctional coupling

*I*_m_ is an integral part of formulas used to define the safety factor of propagation in a one-dimensional tissue (Shaw & Rudy, 1997; Boyle & Vigmond, 2010) and varies as a function of *R*_i_ (**Eq. 5**). The simulations in the previous section for the hiPSC-SM strand were performed with *R*_i_ = 4 MΩ. Shown in **Fig. 15** are additional simulations for *R*_i_ = 20 and 100 MΩ.

As *R*_i_ increases, the propagation delay between adjoining cells increases, *θ* decreases, and the time until cell 35 activates becomes longer (top row, **Fig. 15B**). *I*_in_, *I*_out_ and *I*_m_ (**Eq. 5**) all decrease in amplitude, as does *I*_Na_ (owing to slower rise of *V*_m_ to reach threshold and subsequent greater inactivation of the gating variable *h*), whereas the other ionic currents are relatively unaffected. Although increasing pacing rate and increasing *R*_i_ both slow *θ* (**Figs. 14** and **15)**, increasing pacing rate (CL of 2000 ms to 500 ms) has a much greater shortening effect on AP duration and inhibitory effect on *I*_Na_ (smaller amplitude and greater elongation) than does intracellular resistance increasing from 4 to 200 MΩ.

**Figure 14.**
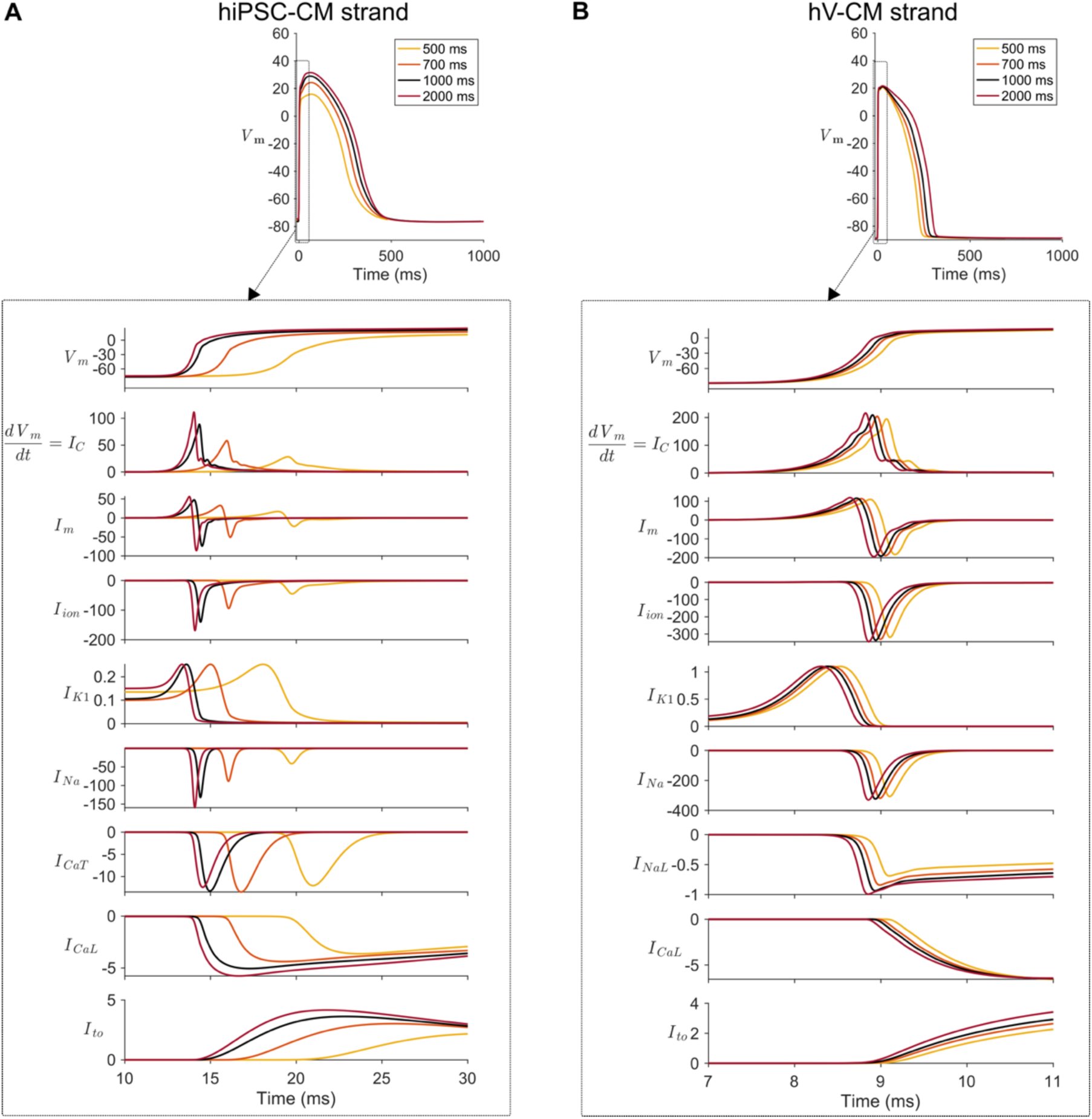
Excitation of cell 35 in a strand at different pacing rates. **A.** hiPSC-CM strand (mKC1 cable) with *R*_i_ = 4 MΩ. **B.** hV-CM strand (ToR-ORd cable) with *R*_i_ = 1.67 MΩ. Plotted in the rows are four sets of *V*_m_ (mV), *dV*_m_/*dt*, and ionic currents, for pacing CLs of 2000 ms, 1000 ms, 700 ms and 500 ms. See caption for **Figure 11** for more details on units. Insets show the full APs at the four pacing rates. Units of *V*_m_ is mV, d*V*m/dt is mV/ms, and current is A/F.

**Figure 15.**
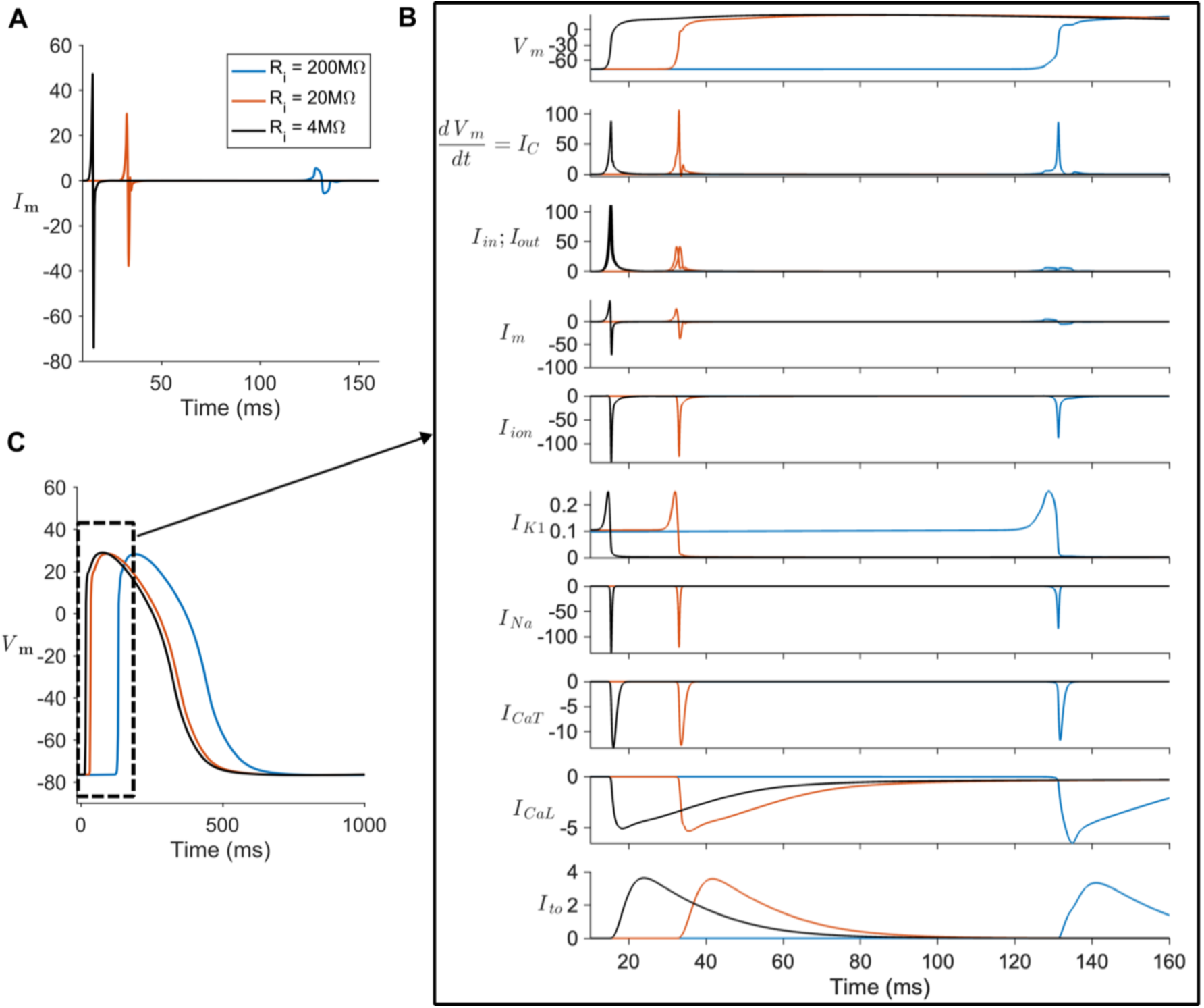
Voltage and ionic currents for cell 35 in strands with differing *R*_i_ in the hiPSC-CM strand. **A.** *I*m. **B.** *V*_m_ and ionic currents. Time 0 is time of stimulation at the initial end of the strand. *R*_i_ = 4, 20 or 200 MΩ. **C. APs** corresponding to the three *R*_i_ values. Pacing CL = 1000 ms. Units of *V*_m_ is mV, d*V*m/dt is mV/ms, and current is A/F.

As in **Fig. 12A**, we approximated *I*_m_ as a biphasic triangular pulse. The amplitude and duration of its two phases along with the interpeak interval are plotted in **Fig. 16** for *R*_i_ ranging from 1 to 1000 MΩ.

**Figure 16.**
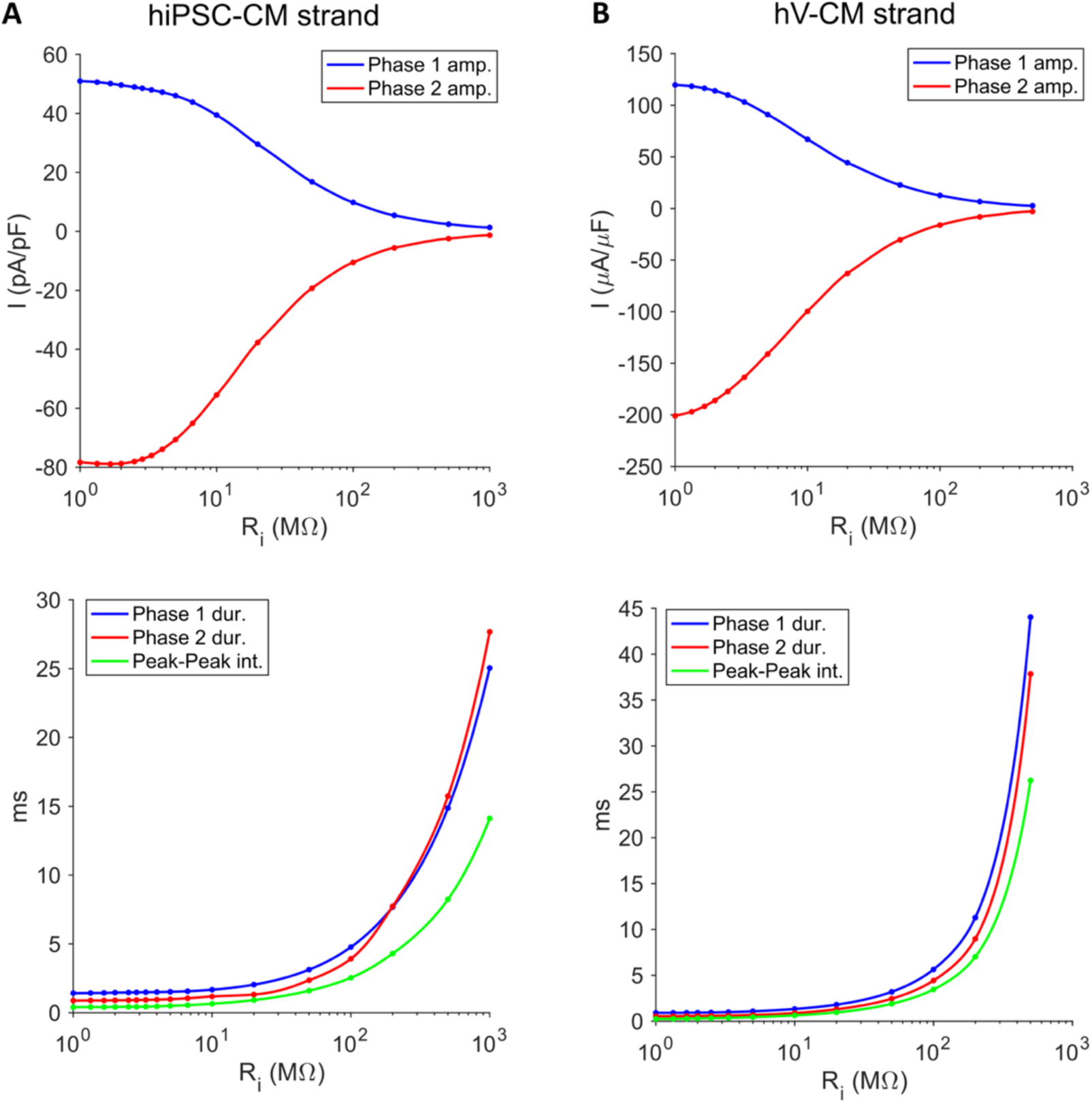
Biphasic triangular approximation to *I*m for different *R*i. **A.** hiPSC-CM strand (mKC1 cable) **B.** hV-CM strand (ToR-ORd cable)

For hiPSC-CMs and hV-CMs, the total duration of the biphasic pulse (sum of durations of the two phases) varies with *R*_i_, and for *R*_i_ between 1 and 10 MΩ, it ranges from 2.3 to 2.9 ms for hiPSC-CMs and from 1.4 to 2.3 ms for hV-CMs. For larger *R*_i_ the total duration can become very long, reaching 52.7 ms (at *R*_i_ = 1000 MΩ) for hiPSC-CMs and 81.9 ms (at *R*_i_ = 500 MΩ) for hV-CMs. Thus, the decrease in *I*_Na_ at high *R*_i_ (> 10 MΩ, **Fig. 15B**) is due to the increase in effective pulse duration of *I*_m_, which allows more inactivation of gating variable *h* to occur, akin to the reduction in *I*_Na_ in single cells with increasing stimulus pulse durations (**Fig. 10**).

### 5. Drug Block

#### 5.1 Population models

For the last part of this study, we examined the effect of varying degrees of Na^+^ channel block in a population of cells having heterogeneous properties in the form of slightly different levels of ion channel expression. This is of practical interest, because hiPSC-CMs grown in culture are a heterogeneous population affected by cell density and culture conditions (Du *et al*., 2015; Li *et al*., 2020). A heterogeneous population can also reveal varying cellular responses to drug block (Sarkar & Sobie, 2011; Muszkiewicz *et al*., 2016; Ni *et al*., 2018). To achieve this population, we varied the maximal conductances of all 16 of the ion channels, transporters and pumps, as well as the baseline values for cell capacitance and cell volume, as normal Gaussian distributions with ±10% standard deviation. These values were then sampled in random combinations to create an initial set of 10,000 variants of both the mKC1 and mKC2 models. Not all the models displayed physiological behavior – for example, for mKC1, approximately 72% lacked spontaneous activity, while 20% failed to repolarize in a normal fashion or were spontaneous and failed to capture.

Hence, these models were filtered out based on the criteria outlined in the **Appendix**. With mKC1 as the baseline model, 1119 models passed the filtration algorithm and of these, 164 satisfied the experimental data at all four pacing rates. With mKC2 as the baseline model, 3108 models passed the filtration algorithm, 492 of which satisfied the experimental data.

We also created a population of models using Latin hypercube sampling (LHS) of up to ±30% variation of the maximal conductances and transport rates of the ion channels, transporters, and pumps, and values of cell capacitance and cell volume. With mKC1 as the baseline model, 6939 out of 60,000 initial models passed our filtration algorithm and of these, 156 satisfied the experimental data at all four pacing rates. With mKC2 as the baseline model, 3476 out of 10,000 initial models passed our filtration algorithm, 435 of which satisfied the experimental data. The results using LHS were very similar to those obtained with random sampling of the normal distributions about the baseline values and are presented here. **Fig. 17** superimposes APs from the population of mKC1 and mKC2 models compared with the APs recorded in vitro, as well as the distribution of their respective APD_30_ and APD_80_ values relative to the constraint space defined by our *in vitro* data.

**Figure 17.**
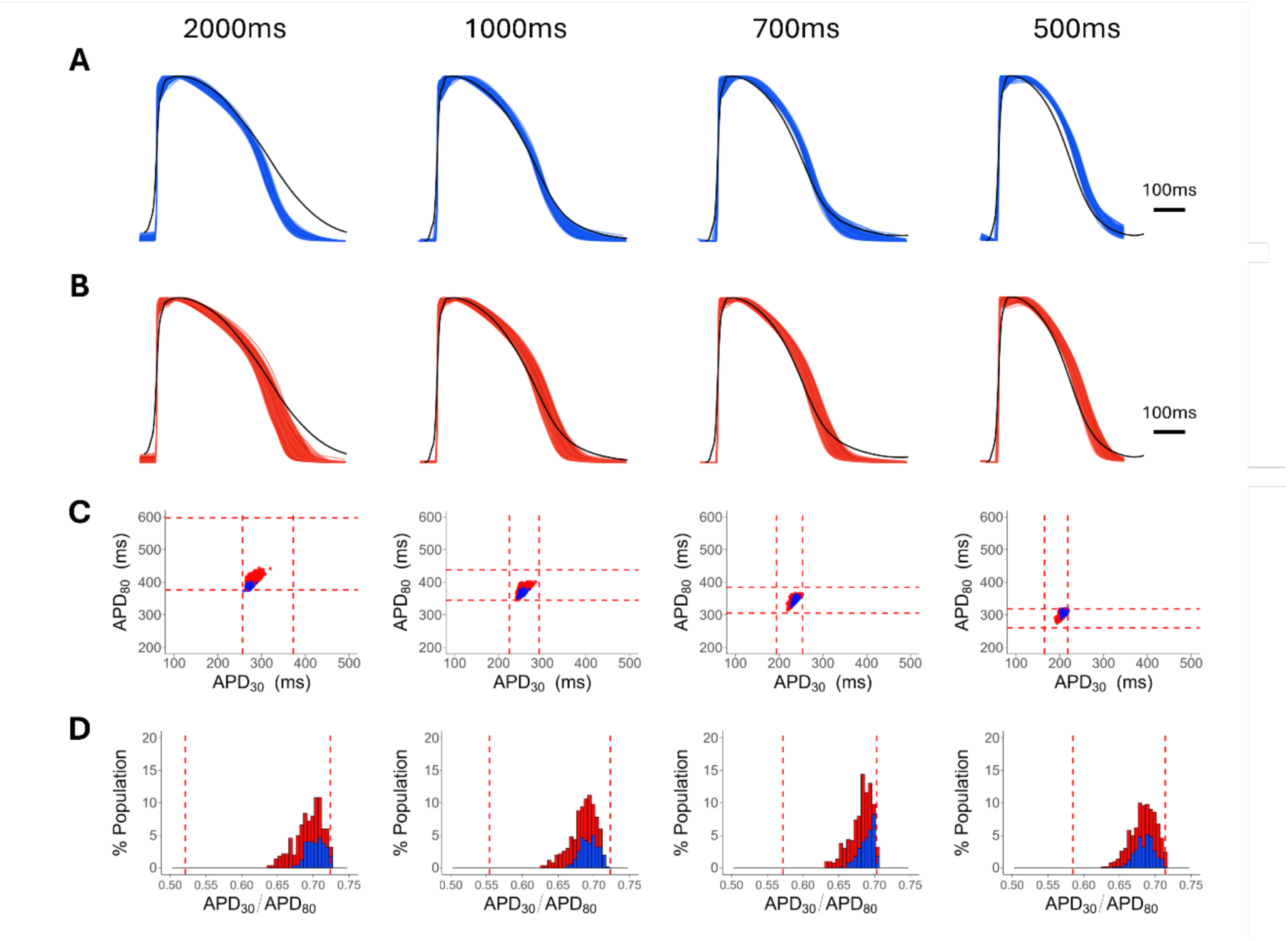
APs and APD measurements for the hiPSC-CM models at different pacing rates. **A.** Superimposed APs for the population of mKC1-based models (n= 156) at different pacing CLs. **B.** Similar plots as in **A** except now for the population of mKC2-based models (n = 435). For both **A** and **B**, the black traces represent the average in vitro AP for the given pacing rate. **C.** Blue points are APD30 and APD80 for each of the mKC1 APs, while the red points are APD30 and APD80 for the mKC2 action potentials. The dashed lines represent the 10% and 90% cutoffs of the experimentally measured histograms of APD30 and APD80 (Table 1). **D.** Histograms representing the APD30:APD80 ratios for the mKC1 (blue) and mKC2 (red) populations.

Similar to **C**, dashed lines represent the 10% and 90% cutoffs from the experimental data. For all rows, columns 1, 2, 3, and 4 show data from 2000ms, 1000ms, 700ms, and 500ms pacing CLs, respectively.

The mKC2 population fills out the experimental constraint space more fully than the mKC1 population, and less scatter is seen with the mKC1 population.

Next, the mKC1 population was stimulated at 1.5x threshold, and *g*_Na_ was reduced by 10%, 50% and 90% (**Fig. 18**). With increasing current block, a greater fraction of the population failed to excite (seen as subthreshold responses).

**Figure 18.**
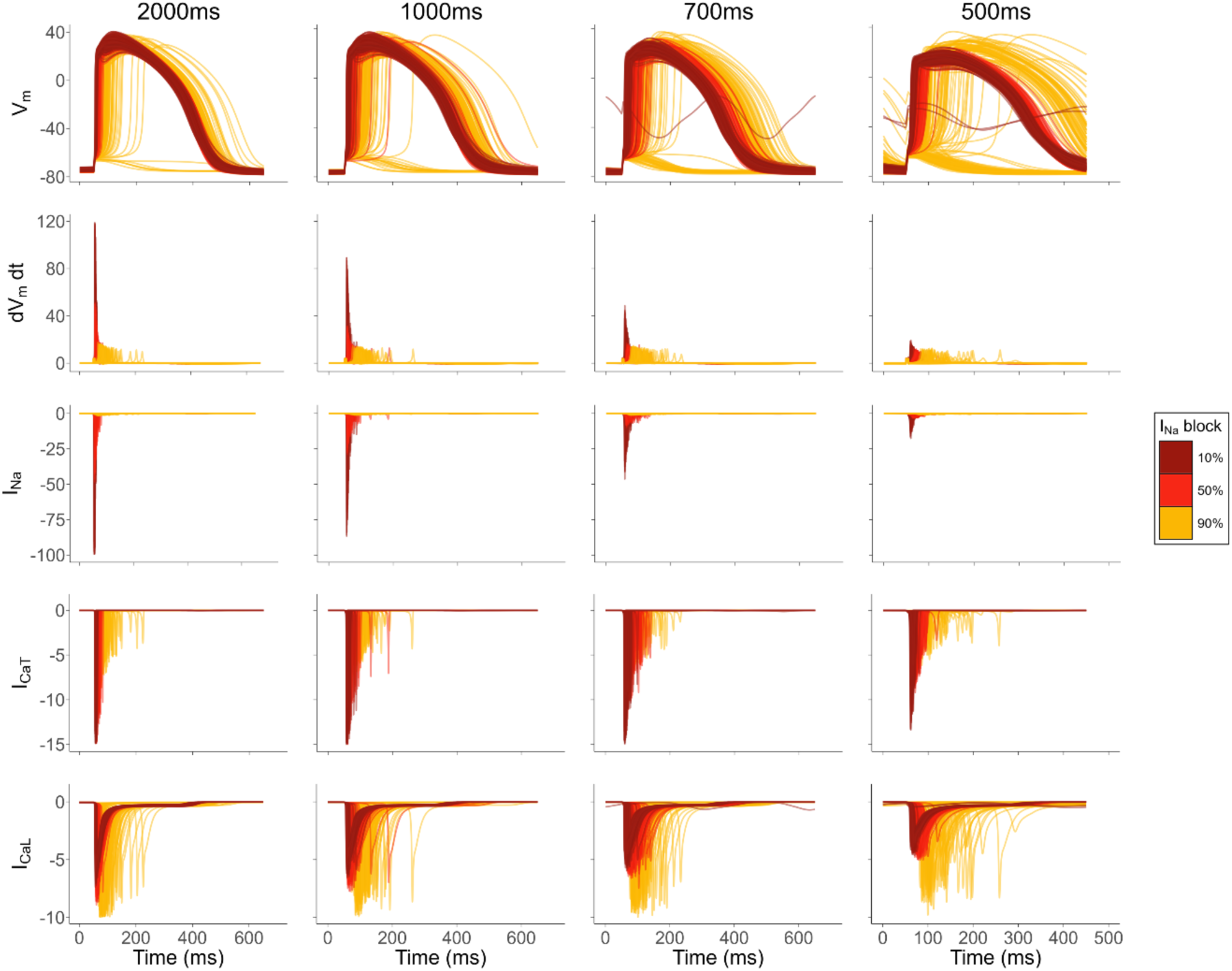
APs and inward ionic currents of models belonging to mKC1 population under varying degrees of *I*Na block at 1.5x stimulus threshold. 156 models of mKC1 population were paced at 2000ms (leftmost column), 1000ms (second column), 700ms (third column) and 500ms (rightmost column) CL. For each pacing rate, after 100 beats all 156 models were subjected to 10% (bright red), 50% (dark red), and 90% (black) *I*Na block. At each pacing rate, the models were stimulated at 1.5x their respective stimulus thresholds. Units of *V*_m_ is mV, d*V*m/dt is mV/ms, and current is A/F.

At only 10% channel block, the entire population could be stimulated at all four pacing rates, although the peak of the AP and magnitude of *I*_Na_ diminish at higher rates. None of the models exhibited repolarization failure, with the exception of one at 700 ms and two at 500 ms CL. However, at 90% channel block, much of the population (96% at 2000ms, 91% at 1000ms, 79% at 700ms, and 44% at 500ms) could still be stimulated. *I*_Na_ is virtually absent, whereas the excitatory inward currents consist mainly of *I*_CaT_ and *I*_CaL_. The ability to excite despite varying levels of *I*_Na_ block is also observed experimentally with iPSC-CMs, as is the increasing delay in upstroke with increasing level of block (Ma *et al*., 2011).

To quantify changes in the composition of the population, *t*_lag_ was used as a measure of the proximity of stimulus strength to *I*_T_ (as seen in **Figs. 3** and **4**), and results are shown in **Fig. 19** for the three levels of channel block at each of the four pacing rates. Only those with activated APs are shown.

**Figure 19.**
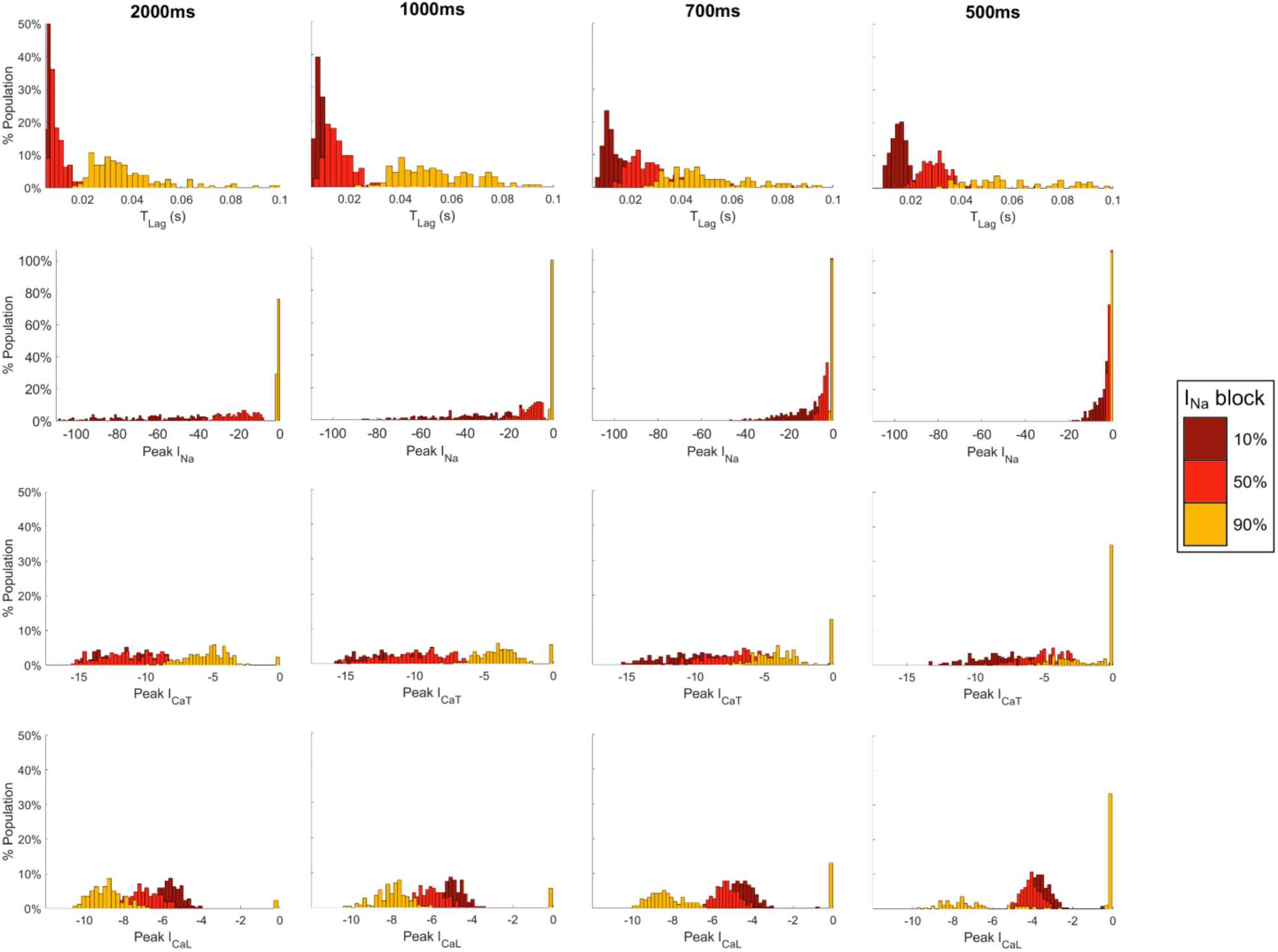
Histograms of *t*lag of mKC1 population under varying degrees of *I*Na block with stimulus strength at 1.5x threshold. The mKC1 population was paced at 2000ms (leftmost column), 1000ms (second column), 700ms (third column) and 500ms (rightmost column) pacing CL. For each model, the lag time from stimulus onset to maximum upstroke (*t*lag) is quantified as well as the peak negative value for each inward current. At each pacing rate, all models were stimulated at 1.5x their respective stimulus thresholds. For clarity, peak *I*Na values are presented as stacked histograms.

At each of the four pacing rates, the distribution of *t*_lag_s beomes longer and wider with increasing block, indicating that *I*_T_ is increasing and becoming more variable across the population. *t*_lag_ is quantified only for models that are able to excite for the given combination of *I*_Na_ block and pacing rate, as *t*_lag_ is undefined for models that fail to excite. However, peak *I*_CaT_ and *I*_CaL_ with values close to zero do indicate non-excited models, as neither current becomes activated.

## Discussion

Experimental studies of cardiac action potentials in whole hearts and tissue explants date back for more that 50 years, with early work summarized in the seminal textbook by Hoffman and Cranefield (Hoffman *et al*., 1960). The development in the 1980s of the single cardiomyocyte under patch clamp conditions opened the door for the biophysical characterization of the ionic currents underlying the action potential that in turn, led to the development of computational models of the cardiac cell, which today underlie many large scale models of the heart and the study of arrhythmogenesis and potential therapeutic strategies. The revolutionary development of cardiomyocytes derived from human induced pluripotent stem cells (hiPSC-CMs) in the late 2000s raise realistic prospects for personalized treatment of genetic disorders and population studies of cardiac diseases.

Multivariable linear regression has been used to determine the sensitives of biomarkers to biophysical parameters (Sobie, 2009) and to predict AP and calcium transient biomarkers in hV-CMs using hiPSC-CM data (Gong & Sobie, 2018), while logistic regression, has been applied to predict arrhythmic events like EADs and to identify underlying factors (Morotti & Grandi, 2017). Population modeling accounts for physiological and pathological variability and when calibrated with experimental data, provides insights into population subgroups that may be particularly susceptible to drug effects or disease conditions as well as ionic mechanisms that may underlie specific cardiac arrhythmias (Muszkiewicz *et al*., 2016; Ni *et al*., 2018). These approaches can help to identify influential parameters and potential therapeutic targets related to cardiac arrhythmia. Nonetheless, evaluating arrhythmia susceptibility can also benefit from detailed analysis of the functional electrophysiology of the cell, including its underlying excitatory properties.

The possibility of arrhythmic activity of hiPSC-CMs is a critical consideration in the application of these cells for cardiac repair. When engrafted, a mismatch of the electrophysiological properties of these cells with those of the host tissue could result in ectopic activity or in conduction block leading to wavebreaks and formation of reentrant activity (circus movement and spiral waves). Thus, a deeper understanding of the processes underlying cellular excitation at both the single cell and tissue levels is valuable to anticipate such problems. Conversely, the absence of arrhythmic activity is key in the screening of new therapies for safety and efficacy. Drug trials in safety pharmacology focus on the QT interval (corresponding to ventricular APD), and in cell assays have centered on EAD formation and irregular beat rate, which represent cell-autonomous mechanisms of focal activity. However, reduction in cellular excitability is also consequential in terms of creating conduction abnormalities that can lead to arrhythmias.

The AP upstroke (phase 0) is synonymous with cardiac excitation, and its associated action currents are the driving force for electrical propagation of the cardiac impulse throughout the myocardium. We were particularly interested in excitation not only well above threshold but also just at threshold, because in this instance electrical propagation is most susceptible to failure, leading to conduction block. In this study we used current, state-of-the-art computational models of the hiPSC-CM and hV-CM to probe the biophysical processes underlying cardiac excitation, and to contrast differences in the excitation of the two cell types which lie at different ends of the developmental spectrum. In addition, we sought to contrast the differences in excitation of the two cell types both as single cells and as cells within a rudimentary tissue (1D strand) containing intercellular interactions and electrical loading effects.

A significant challenge in this study was adjustment of the hiPSC-CM model to reflect the AP shape, which in our lab and others (Paci *et al*., 2013; Gunawan *et al*., 2021; Seibertz *et al*., 2023) often have a slow repolarization tail. This could reflect the immature state or phenotype of the cells, since it is absent in adult ventricular cardiomyocytes. Existing models of *V*_m_ of hiPSC-CMs (Koivumäki *et al*., 2018; Kernik *et al*., 2019; Paci *et al*., 2020) lack such a slow repolarization tail. A recent study from the Sobie lab (Yang *et al*., 2024) used a genetic algorithm to optimize the fit of the KC model to optical recordings of hiPSC-CM APs, which resulted in a different set of model parameters than those we have used here. Their model, too, lacked the slow tail of repolarization. The simplest parameter change in the KC model that we could make which prolonged the repolarization tail was an increase in the *α* component of the *I*_Kr_ activation gating parameter, which effectively scaled down the time constant of the gating parameter. However, this increase also weakened repolarization. Although not presented, we found that reduction of *I*_Kr_ produced repolarization failure in a majority of the population of mKC1 cells with 10% block and in a majority of the population of mKC2 cells with 20% block. Thus, the biophysical basis of the repolarization tail has yet to be fully elucidated.

The property of electrophysiological refractoriness, which closely follows APD, affects excitation at rapid pacing rates where successive APs impinge on one another, such that the excitability of an AP is diminished by the preceding AP. The rate-adaptation of APD is well known in iPSC-CMs (He *et al*., 2003; Zhang *et al*., 2009) and is reflected in our study (**Table 1**). Differences between APD_30,_ APD_80_, and the ratio APD_30_/APD_80_ of the original KC model and those of our *in vitro* data at four different pacing rates warranted further modifications to the computational models. In the mKC1 model, APD_30_ and APD_80_ were brought in line by decreasing the permeability of *I*_CaL,_ decreasing *I*_f_ and increasing *I*_K1_ slightly, although the rate-dependence of APD remained weaker than that observed experimentally (**Appendix Fig. S1**). In the mKC2 model, the slope of the rate-dependence was increased by a synergistic combination of a reduction in sarcolemmal pump activity, a negative voltage shift of the *I*_Ks_ activation parameter, a reduction of the *I*_CaL_ inactivation time constant, and an increase in *I*_K1_ conductance, but was still weaker than that observed experimentally. This weakness is shared among other computational CM models (**Appendix Fig. S1**).

The major excitatory currents during the AP upstroke are *I*_Na_, *I*_CaT_ and *I*_CaL_ in the mKC1 model of the hiPSC-CM, and *I*_Na_, *I*_NaL_ and *I*_CaL_ in the ToR-ORd model of the hV-CM. Depending on the strength of the stimulus above threshold and its duration, the delay in the upstroke and the magnitudes of the ionic currents can vary. This finding is important, because electrophysiological studies which measure or model APs and ionic currents in single cells often do not report the stimulus waveform’s shape, amplitude, and duration. The situation for tissue is different, however, since the stimulus current is not under external control but rather, is determined by the excitatory wavefront. When the stimulus is increased to twice threshold, *t*_lag_ becomes shorter, and in hiPSC-CMs, *I*_Na_ and *I*_CaT_ increase while *I*_CaL_ decreases (**Fig. 4**). The peak values of the other ionic currents are essentially unchanged. Like in hiPSC-CMs, in hV-CMs, *I*_Na_ also increases, and *I*_CaL_ decreases. However, the peak values of the outward currents *I*_to_, *I*_NaK_, *I*_K1_ and *I*_NaCa_ are more positive, due to the larger overshoot of the AP, which arises because AP activation is so fast that it occurs within the duration of the stimulating pulse.

Excitability of cells and tissues at rest is often characterized by the strength-duration relation, which maps out the minimum stimulus strength needed to excite the cell for varying pulse durations (**Fig. 8**). In general, the SD relations to excite the end of a strand are elevated above those of the single cell (for both hiPSC-CMs and hV-CMs), because of the loading effect of downstream passive cells. In neonatal rat cell cultures, the SD relation for 3D aggregates is elevated compared with that for 2D monolayers (Sayegh *et al*., 2019) and may also be a manifestation of the greater loading effect of downstream cells. In comparing hiPSC-CMs to hV-CMs, hiPSC-CMs are more excitable, as seen in their lower SD relations for both single cells and strands. It is also reflected in subthreshold stimulation, in which decremental conduction extends for longer distances (**Fig. 5**). A reason for the greater excitability of hiPSC-CMs is the intrinsic automaticity of the cells, in which inward currents (largely *I*_f_, absent in hV-CMs) are actively working against the background outward currents (primarily *I*_K1_ which is weakly expressed in these cells) to depolarize the cell.

Furthermore, the more positive *V*_m0_ of hiPSC-CMs is closer to their *V*_T_ (a difference of about 7 mV at 1000 ms pacing CL) compared with the difference in these potentials in hV-CMs (about 25 mV at the same pacing CL). When engrafted, the mismatch of excitability of hiPSC-CMs with that of the host tissue could result in ectopic activity that could trigger premature ventricular contractions or even tachyarrhythmias. Hence, our study presents a cautionary note for applications using these cells, particularly for cardiac repair.

SD relations are typically characterized by two parameters, their rheobase and chronaxie (**Eq. 3**). The computational models used in this study allow extremely long pulses to be applied which are not attainable in practice. These result in very low values of rheobase (and subsequently, large values of chronaxie) compared with rheobases defined at an ad hoc pulse duration (such as in (Kay *et al*., 1990) or **Table 4**). The values for chronaxie for hV-CMs (based on the ToR-ORd model) are much greater than those reported in the literature, which are typically less than 1 ms (Kay *et al*., 1990; Geddes, 2004; Jastrzębski *et al*., 2019). Another contributing factor for the discrepancy may be that in practice, the stimulation of cardiac tissue at the organ level is typically applied extracellularly as opposed to intracellularly as in this study. Extracellular stimulation involves a fundamentally different biophysical process (Sobie *et al*., 1997) that can give rise to spatially complex patterns of cellular polarization (Basser & Roth, 2000) that could affect the measurement of rheobase and consequently, chronaxie. In the design of pacemakers, the rheobase carries significance in terms of the minimal current that is required of the device to stimulate (although applicable only at long pulse durations) as well as the battery requirements. Chronaxie is, by some theoretical calculations, most energy efficient for stimulation (Geddes & Bourland, 1985) and has been used as a guide for the optimal pulse duration (Coates & Thwaites, 2000). The rheobase is strongly influenced by *I*_K1_ conductance in both cell types, which is in accord with the findings of Hund and Rudy who showed that excitability is governed by membrane resistance for long duration stimuli (Hund & Rudy, 2000). We also found the rheobase to be highly sensitive to the pump current *I*_NaK_ in hiPSC-CMs and, to a smaller extent, in hV-CMs (**Fig. 9**). The chronaxie is also highly sensitive to *I*_K1_ conductance in both cell types, and to a lesser degree on *I*_NaK_ in hV-CMs (**Fig. 9**). However, the dependence of chronaxie on other ionic currents is not straightforward. We also fit the SD relations with single and double exponentials to determine whether any of the parameters in those equations were strongly related to individual ionic currents (not shown) but were unsuccessful in this regard.

The relative amplitudes of ionic currents underlying cell excitation vary somewhat at different stimulus pulse durations when stimulating at twice threshold. As pulse duration increases from 1 to 20 ms, the most notable change is a decrease in *I*_Na_ for hiPSC-CM and decrease in *I*_CaL_ for hV-CM (**Fig. 10**). These decreases may be attributed to the substantial delay after the end of the stimulus pulse for *V*_m_ to reach *V*_T_ (*t*_lag_, which is relatively independent of rate, **Table 3**), during which time some inactivation of the currents can occur, more so for *I*_Na_ in hiPSC-CMs and for *I*_CaL_ in hV-CMs. Although we did not study how different durations of conditioning trains effect excitability, the dependence of excitability on pacing history (memory effect) was extensively studied by Hund and Rudy and shown to be mediated by changes in the membrane resting resistance *R_m_* for long duration stimulus pulses, secondary to changes in intracellular concentrations of Na^+^ and Ca^2+^ (Hund & Rudy, 2000).

Previous studies have examined the success or failure of conduction in cardiac tissue in terms of safety factor and have identified *I*_Na_ and *R*_i_ as two key parameters (Shaw & Rudy, 1997; Boyle *et al*., 2019). Our study further clarifies the excitation of hiPSC-CMs by a propagating AP (in a strand) in comparison with that of an isolated cell stimulated by an external current pulse (**Figs. 11** and **12**). The portion of *I*_in_ that does not leave the embedded cell via *I*_out_ is the cell’s stimulus current, *I*_m_. Unlike the monophasic rectangular pulse used to calculate the strength-duration relation, *I*_m_ during excitation (AP activation) is biphasic and more triangular shaped. For an intracellular resistance of 4 MΩ, we showed that *I*_m_ can be closely approximated by a biphasic triangular pulse. Thus, while the strength-duration relation for monophasic pulses is relevant for electrical stimulation of cardiac tissue, it is not directly applicable for physiological stimulation by a propagating AP. Notably, Shaw and Rudy (Shaw & Rudy, 1997) used only the first positive phase of *I*_m_ in their definition of safety factor. Our results demonstrate that the second, negative phase of *I*_m_ acts to boost the amount of membrane ionic current (*I*_ion_) available as the source current to stimulate downstream cells.

*I*_in_ is the electrical source current provided to the cell by its upstream neighbor, or conversely, the load that the cell places on its upstream neighbor. Until *V*_m_ of the cell reaches *V*_T_, *I*_in_ is primarily *I*_C_ (with a small contribution from *I*_K1_). Once *V*_m_ reaches threshold, *I*_Na_ is activated and takes over the charging of membrane capacitance, and *I*_in_ declines as the electrical load to the upstream neighbor dissipates with cell activation.

Meanwhile, *I*_out_ is the electrical current drained from the cell by its downstream neighbor, or conversely, the current that the cell provides to excite its downstream neighbor. *I*_out_ includes a portion of *I*_in_ from the upstream neighbor, which constitutes an electrotonic effect extending to the downstream cells. Once the cell reaches threshold, *I*_Na_ is activated and now takes over as the primary component of *I*_out_. Although it reaches a peak and decays, *I*_out_ continues to be supplied via *I*_CaT_ and to a lesser extent by *I*_CaL_. Together, *I*_Na_, *I*_CaT_, and *I*_CaL_ act as current sources which flow from the cell to depolarize its downstream neighbor up to *V*_T_ and subsequently trigger an AP. *I*_out_ then declines as the electrical load from the downstream neighbor dissipates with its activation.

Cell excitability affects the conduction velocity of electrical wavefronts. As we have shown, increasing pacing rate depresses excitability by reducing excitatory currents, raising *I*_T_ (**Table 4**), and slowing *θ* (**Appendix Table S1**). *V*_T_ does not vary much, at least in normal cells, and is not a sensitive index (**Table 3**). Excitability is also affected by the amount of electrical loading from neighboring cells. Thus, if the strength of stimulation were to hover near threshold, small differences in electrical loading at bifurcations (branched networks) could lead to failure in one branch and not the other, potentially leading to a reentrant pathway with unidirectional block. Even with equivalent loading (*R*_i_), small differences in the levels of excitatory currents could lead to conduction failure in one branch and not the other.

An increase in intracellular axial resistance (e.g., from a decrease in gap junctional coupling) results in diminished *I*_in_, *I*_out_ and *I*_m_. This necessitates a prolongation of these currents to bring *V*_m_ up to *V*_T_ (**Fig. 15**). However, this prolongation also allows for some inactivation of the excitatory currents to occur, and in this sense, it reflects the strength-duration curve at long pulse durations. Indeed, we found a greater degree of inactivation of *I*_Na_ with larger *R*_i_, which reduces *I*_ion_ and in turn, reduces *I*_C_ (and thus, (*dV*_m_/*dt*)_max_), as well as *I*_out_, the source current for downstream cells. Despite the reduction in *I*_Na_ with increasing *R*_i_, the safety factor for propagation can increase (up to a point) (Shaw & Rudy, 1997) because less *I*_Na_ flows into downstream cells and more into *I*_C_. Excitation of downstream cells can still occur so long as their *V*_m_ reaches threshold and their own inward currents, *I*_Na_, *I*_CaT_, and *I*_CaL_ can excite the cell. Another consequence of larger *R*_i_ is that because *I*_m_ is prolonged, there is now sufficient time during its initial positive phase for *I*_Na_ (and *I*_CaT_ for hiPSC-CMs) to inactivate such that its negative phase does not enhance these currents, although *I*_CaL_ and *I*_to_ (and *I*_NaL_ for hV-CMs) are still affected.

Finally, our population models with both the mCK1 and mKC2 formulations demonstrate that sodium channel (*I*_Na_) block can delay AP activation by varying degrees, which increase as the level of channel block or beat rate increases, or as the stimulus strength decreases (**Figs. 18-20**). Some cells remain excitable even at the fastest beat rate and higher levels of sodium channel block, while others fail almost immediately. The percentage of such excitation failures increases in a rate-dependent manner (**Fig. 20**). Such behavior can potentially induce regional block and reentry circuit behavior in tissue containing grafts of hiPSC-CMs having such heterogeneity and represents a proarrhythmic mechanism distinct from cell-autonomous arrhythmias arising for example, from early afterdepolarizations.

## Limitations of the Study

The form of stimulation used in this, and many other studies is current injection into the cell or strand. The classical strength-duration relation is based on monophasic pulses, whereas physiological stimulation involves biphasic pulses. However, stimulation by extracellular electric fields, such as with pacing electrodes, involves a biophysical process distinct from current injection, that produces coexisting regions of depolarized and hyperpolarized cell membrane (Sobie *et al*., 1997; Tung & Kléber, 2000; Sharma & Tung, 2002). These regions can result in both cathodal and anodal excitation following the make or break of the stimulus pulse (Basser & Roth, 2000). Computational simulations would need to take this complexity into account.

To accommodate observations of a slow tail of repolarization in our hiPSC-CMs, we altered one of the kinetic parameters of the model, but this also weakened repolarization. Although repolarization per se was not the subject of this investigation, its kinetics presumably become important at fast pacing rates, and therefore error in its representation could be a limitation of our study. However, the majority of our simulations were performed at a 1000 ms pacing CL (1 Hz rate), where such effects may be minor.

To study intercellular source-load effects, we simulated a strand of cells as a 1D cable. To reduce computational expense, we limited the length to 50 cells and the runtime to 100 beats of conditioning prior to the test stimulus. The cable also consisted of discrete cell membranes separated by intracellular axial resistances, which did not allow for spatial gradients of *V*_m_ within individual cells or distinguish between myoplasmic and gap junctional resistances. However, the error between this kind of model and models containing subcellular compartments and separation of myoplasmic and gap junctional resistances may be small (Shaw & Rudy, 1997). A recent study by Pullinger and Sobie examined the vulnerability to arrhythmia of a 1D cable containing a heterogenous population of ToR-ORd cells (Pullinger & Sobie, 2024).

In our simulations of the dependence of excitability on *R*_i_, we assumed that *I*_Na_ was unchanged. However, this is not necessarily the case. Overexpression of Cx43, which would be expected to increase gap junctional conductance and decrease *R*_i_, can result in a gain of function of the fast Na channel, leading to enhanced *I*_Na_, AP upstroke velocity and presumably, excitability (Sottas *et al*., 2018).

Another limitation of our study is the assumption of electrical conduction via ionic currents passing through intracellular resistive pathways containing gap junctions. Experimental studies have shown that even with genetic knockout of connexin43 (the principal gap junction protein in the ventricle), electrical propagation persists although with a higher incidence of arrhythmias (Gutstein *et al*., 2001). These findings underscore the necessity to include other forms of intercellular electrical connections such as ephaptic coupling (Weinberg, 2017; Lin *et al*., 2022), particularly at low levels of gap junctional conductance.

In our simulations of *I*_Na_ block, the block was constant across pacing rates, whereas in reality, it can increase at faster pacing rates due to more frequent exposure of the drug to binding sites within the sodium channel (Starmer *et al*., 1984). More generally, the drug block of other ion channels can be rate-dependent, with block either increasing at higher pacing rates, or increasing at lower pacing rates (reverse rate-dependence), the latter of which is a proarrhythmic factor (Shah & Hondeghem, 2005). Such dependencies could be incorporated in future studies.

Lastly, the immature state of hiPSC-CMs is also an important factor influencing excitability, as we have shown in comparison with hV-CMs. hiPSC-CMs are in a state of flux as they mature, with expression of Na and other channels increasing with time (potentially increasing excitability) and automatic activity decreasing with time (potentially decreasing excitability). Furthermore, electrical pacing itself can accelerate the maturation of hiPSC-CMs (Ronaldson-Bouchard *et al*., 2018) and act as positive feedback to increase excitability. We also assumed a specific size of cell and density of cells in our model of a 1D strand.

However, it is known that these parameters, as well as cell orientation, presence of other cell types, metabolic state, and disease conditions can affect excitability (Tung *et al*., 1991; Du *et al*., 2015; Sayegh *et al*., 2019; Li *et al*., 2020). These factors can be engineered into *in vitro* systems and simulated computationally. Further, cells grown as higher order dimensional tissue (e.g., 3D engineered heart tissues) can have greater Na channel expression and enhanced excitability not accounted for models based on single, isolated cell measurements (Lemoine *et al*., 2017).

## Conclusions

Computational models of hiPSC-CMs were generated that complied with experimental measurements in our laboratory of APD_30_, APD_80_ and their ratio at four pacing rates. The excitation of hiPSC-CMs at and above the threshold of stimulating APs was compared with that of hV-CMs, including the involvement of specific ionic currents, the strength-duration relation, and cells embedded within a strand of tissue. In general, hiPSC-CMs were found to be more excitable than hV-CMs in terms of lower stimulus threshold and greater decremental subthreshold conduction, attributable to their intrinsic automaticity.

However, conduction velocity was slower, due to smaller cell size, smaller *I*_Na_, and lower gap junctional conductance. When stimulated at or above threshold, hiPSC-CMs had longer lag times to activation than did hV-CMs. These lag times could be used as a sensitive measure of the proximity of the stimulus amplitude to its stimulus threshold. When stimulating at twice threshold vs. at threshold, peak *I*_Na_ was larger, but peak *I*_CaL_ was smaller for both cell types.

In comparing strands to cells, the stimulus threshold of strands at their ends was higher, due to the loading of downstream cells. For cells in the middle of the strand that were excited by a propagating AP, the physiological stimulus current *I*_m_ is biphasic. Our modeling results showed that *I*_m_ could be closely approximated by a biphasic triangular pulse with a total duration of ∼ 2-3 ms for both cell types (slightly shorter for hV-CMs) for intracellular axial resistances in the range of 1-10 MΩ. Increasing the intracellular resistance increases the stimulus threshold for either cell type in a strand due to the increase in lag time for *V*_m_ to reach threshold, during which time inactivation of *I*_Na_ can occur.

The excitability of both cell types, whether in isolation or embedded within a strand, could be quantified as stimulus threshold (*I*_T_), strength-duration relation, or lag time to activation (*t*_lag_). Excitability was greater in hiPSC-CMs than in hV-CMs and attributed to the intrinsic pacemaking activity of the cell working in the background. Excitability of both cell types decreased in single cells stimulated with longer duration pulses, in single cells and strands with faster pacing rates, and in strands with greater intracellular resistance. The reductions in excitability were generally related to decreased amplitude and elongation of *I*_Na_, arising from larger degrees of inactivation.

Finally, we showed that a given level of sodium channel block across a population of either hiPSC-CMs or hV-CMs results in an increased level and broader dispersion of their stimulus thresholds, together with an increasing fraction of unexcitable cells. This loss of collective excitability was rate-dependent.

## Glossary

AP: action potential
APD: action potential duration
CL: cycle length
CM: cardiomyocyte
*d*: duration of stimulus pulse
(*dV*_m_/*dt*)_max_: maximum rate of rise (upstroke velocity) of *V*_m_ during phase 0 of the AP
hiPSC-CM: human iPSC-derived cardiomyocyte
hV-CM: human adult ventricular cardiomyocyte
iPSC: induced pluripotent stem cell
*I*_T_: stimulus threshold. For the single cell it is the minimum stimulus strength to depolarize the cell to 0 mV, and for the 1D strand, it is the minimum stimulus strength for which an AP can propagate to the last cell of the strand.
KC: Kernik-Clancy formulation
mKC: modified KC
ORd: O’Hara – Rudy formulation
*R*_myo_: myoplasmic axial resistance
*R*_gap_: gap junctional axial resistance
*R*_i_: intracellular axial resistance (*R*_myo_ + *R*_gap_)
*V*_rest_: resting membrane potential
SD: strength-duration
ToR-ORd: Tomek-Rodriguez revision of ORd
*t*_lag_: lag time from stimulus onset to time of (*dV*_m_/*dt*)_max_
*V*_m_: transmembrane potential
*V*_m0_: transmembrane potential at the time stimulus is applied
*V*_T_: the threshold value of *V_m_* above which the upstroke of the action potential is activated (*dV*_m_/*dt* > 0) and below which passive decay to rest occurs (*dV*_m_/*dt* < 0). It reflects the activation threshold for the cell and can vary with the stimulus conditions.
*θ*: conduction velocity

## Appendix

### 1. Derivation of Modified Kernik-Clancy Model (mKC) from Baseline Kernik-Clancy Model

Using the mKC1 and mKC2 models as described in **Table 2**, a starting pool of 60,000 and 10,000 candidate models is generated by varying the conductances of all 16 currents within the original KC model using a Latin Hypercube Sample and scaling the conductance values by up to ±30% of their baseline values. Cell size and capacitance are also similarly varied. The pool of initial models is then passed through a filtration algorithm to identify and remove problematic models. First, a model is run under non-paced conditions and evaluated on its ability to autonomously generate APs. The model is then tested under paced conditions with all our custom written functions (i.e. a function to characterize a single AP) to ensure that it does not result in any runtime errors. Next, the model is evaluated on its ability to reach steady state within 100s and to demonstrate 1:1 stimulus-capture at the given pacing rate. Finally, a stimulus threshold search function is applied to ensure that a non-zero threshold can be found for the given model at the given pacing rate. If the search function executes without any errors, the model is run to steady state using a high stimulus strength of 5pA/pF, and the stimulus threshold is calculated using the conditions at which steady state is reached. Steady state is defined as the run time required for a successive APs of a given model to differ by less than 10% RMS value. These tests are all applied at 2000ms, 1000ms, 700ms, and 500ms pacing CLs. If the given model passes all the tests, it is kept for further evaluation; otherwise, it is thrown out.

After the initial pool of candidate models is filtered, experimental constraints are applied at all four pacing rates. A model is paced at twice its stimulus threshold value for a given pacing rate using 5 ms rectangular current pulses and is evaluated on whether it falls within the APD_30_ and APD_80_ constraints for that pacing rate. The model is also evaluated on whether its ratio of APD_30_/APD_80_ (a measure of AP shape) falls within the corresponding experimental constraints. Models that pass the in vitro constraints at all four pacing rates are passed through stimulus misalign and repolarization failure filters as a precautionary measure in the event that the filtration algorithm is not able to fully filter out all non-functional models.

### 2. Differentiation of Cardiomyocytes from iPSCs

WTC11 iPSCs were cultured and differentiated into iPSC-CMs as previously described (Hawthorne *et al*., 2021). Briefly, on day of differentiation (DD) 9, cultures were switched to RPMI/B27 with insulin and replated on DD10 using RPMI/B27 without insulin supplemented with 10µM Y-27632. The culture medium was changed to just RPMI/B27 with insulin on DD11. Metabolic cardiomyocyte purification was performed from DD12-16, after which iPSC-CM enriched cultures were switched back to RPMI/B27 with insulin.

Cultures were maintained by replacing the culture medium every 48 hours until DD20, at which point cardiomyocytes were seeded onto Geltrex-coated, 10mm diameter Thermanox coverslips at a density of 300,000 cells/cm^2^ and cultured in RPMI/B27 with insulin for an additional 8-10 days prior to experimentation.

### 3. Optical Mapping of iPSC-derived Cardiomyocytes

Optical mapping was performed as previously described (Hawthorne *et al*., 2021). Briefly, monolayers of iPSC-cardiomyocytes were incubated with 10µM blebbistatin to inhibit contraction and with 10µM di-4-ANEPPS, a membrane potential sensitive dye, and Pluronic F-127 for 10 minutes. Cell confluency was verified using brightfield microscopy. Pulse trains with 2000, 1000, 700, and 500ms CLs were generated using a computer-controlled signal generator and applied via parallel line platinum electrodes via an optical isolation unit at an amplitude of 20V and a pulse duration of 10ms. Changes in optical transmembrane voltage across the monolayer were captured using a 100x100 pixel CMOS camera (MiCAM Ultima-L, SciMedia) at a sampling rate of 1000 frames/s and a field of view of 10mm x 10mm. Images were then processed using a custom MATLAB script.

**Table A1.** Computed values of conduction velocities *0* (cm/s) for hiPSC-CM and hV-CM strands at four different pacing cycle lengths. hiPSC-CM strand consisted of mKC1 model cells and *R*_i_ = 4 MΩ. hV-CM strand consisted of ToR-ORd model cells and *R*_i_ = 1.67 MΩ.

|  | 500 ms | 700 ms | 1000 ms | 2000 ms |
| --- | --- | --- | --- | --- |
| hiPSC-CM strand | 5.5 | 7.1 | 8.2 | 9.1 |
| hV-CM strand | 47.2 | 48.0 | 48.4 | 49.1 |

**Table A2.** Rheobase and chronaxie determined from SD relations (Fig. 8). **A.** Values using **Eqs. 3a** and **3b** to fit the data. **B.** Values using **Eqs. 3c** and **3d** to fit the data. RMSE is the root mean squared error of the equation fit.

| A from Eqs. 3a and 3b |  |  |  | B from Eqs. 3c and 3d |  |  |  |
| --- | --- | --- | --- | --- | --- | --- | --- |
| | $I_{rh}$<br>(A/F) | $c$<br>(ms) | RMSE | | $I_{rh}$<br>(A/F) | $c$<br>(ms) | RMSE |
| Single hiPSC-CM | 0.04 | 173.90 | 0.062 | Single hiPSC-CM | 0.08 | 63.94 | 0.065 |
| hiPSC-CM strand | 0.44 | 42.98 | 0.580 | hiPSC-CM strand | 0.73 | 18.41 | 0.728 |
| Single hV-CM | 0.91 | 27.08 | 0.137 | Single hV-CM | 1.42 | 12.54 | 0.308 |
| hV-CM strand | 3.65 | 9.21 | 0.502 | hV-CM strand | 4.75 | 5.87 | 2.681 |

**Figure A1.**
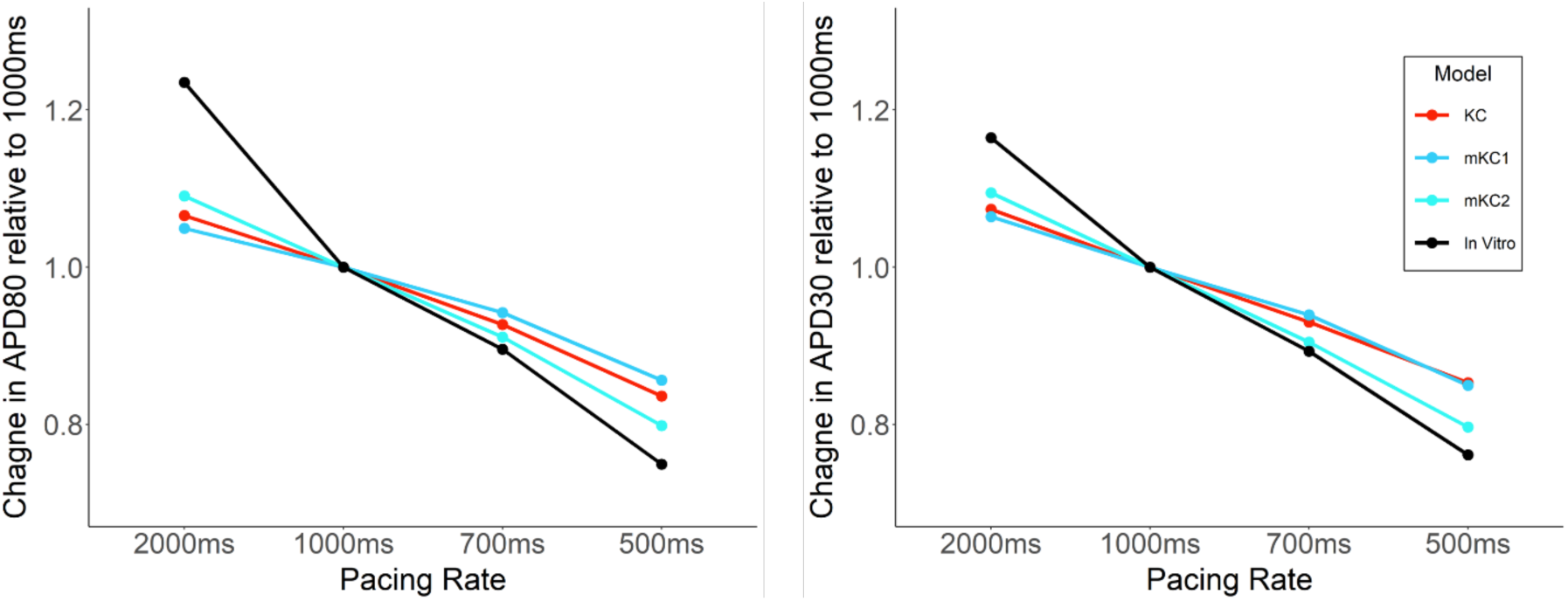
Rate-dependent behavior of APD80 for different computational models. APD80 (left) and APD30 (right) are plotted for the KC, mKC1, mKC2, models and for the in vitro data at different pacing cycle lengths. For each dataset, values were normalized to their values at 1000 ms pacing CL.

**Figure A2.**
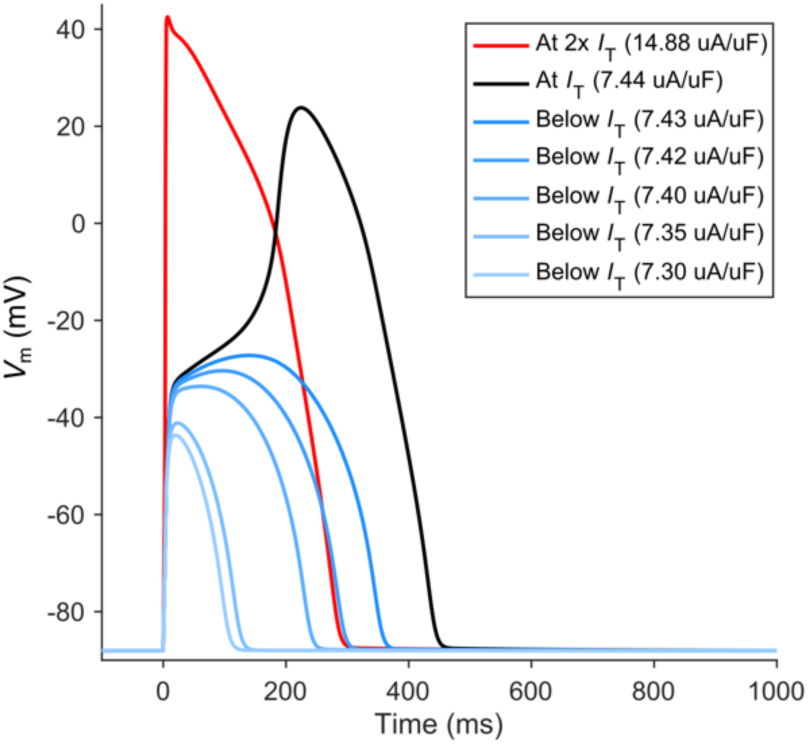
ORd single cell excitation. Stimulus strength varied from just below threshold (blue traces), to threshold (black trace), to roughly twice threshold (red trace).

**Figure A3.**
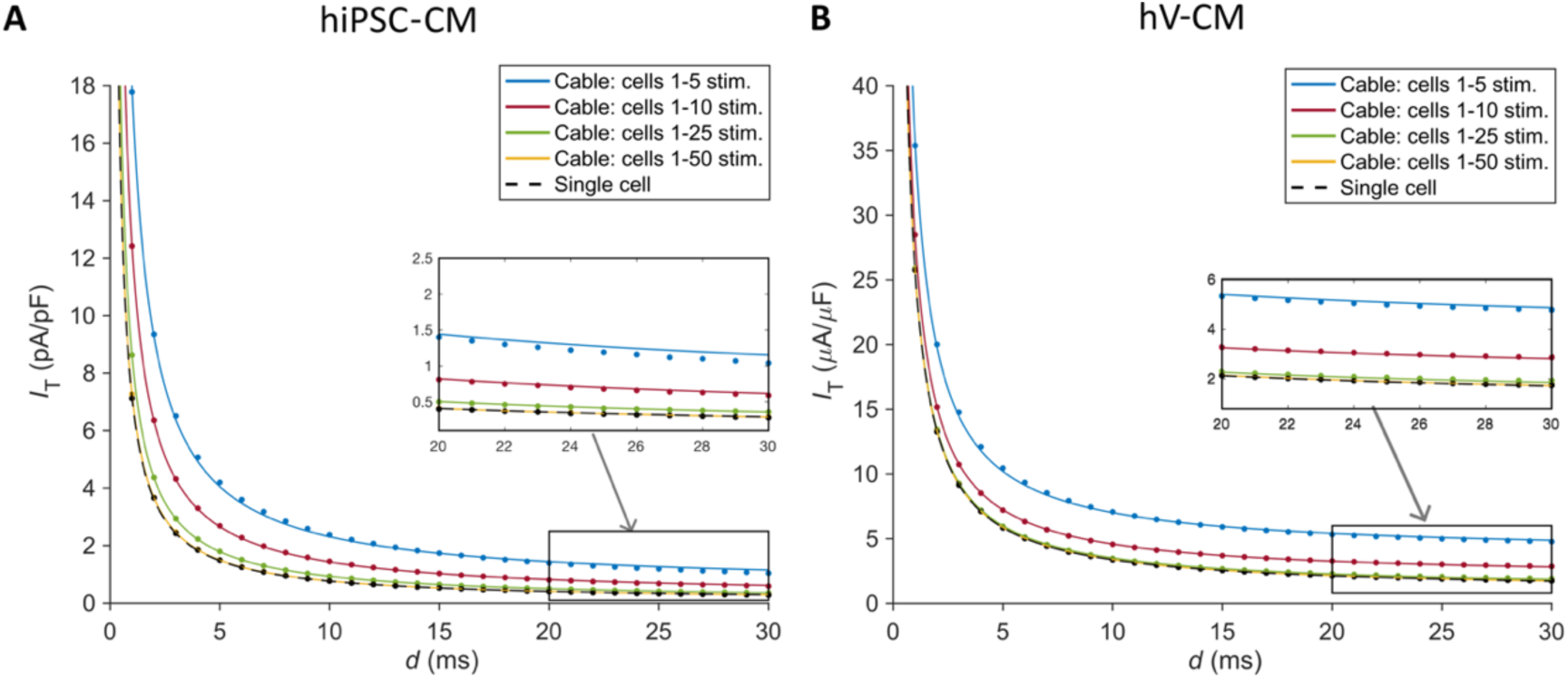
Strength-duration relations for the cable for different numbers of source cells. The number of source cells varies between 1 and 50. Also shown in the black curve is the SD relation for the single cell. **A.** hiPSC-CM (mKC1) model. **B.** hV-CM (ToR-ORd) model. Curves are direct fits using **Eq. 3a**.

## Additional Information

### Data Availability

All source code and instructions will be made freely available on GitHub - https://github.com/HopkinsCBSLab/iPSC_Computational_Excitability

### Author Contributions

RoS and RaS designed and performed the simulations, analyzed the data, and prepared the results, figures and tables. LT conceptualized and supervised the execution of the project, guided the content of the figures and tables, performed some simulations, and provided financial support. All authors participated in the discussion and interpretation of the data. Writing and editing of the manuscript was performed by LT with assistance from RoS and RaS. All authors have approved the final version of the manuscript and agree to be accountable for all aspects of the work.

## Acknowledgements

We are grateful to Dr. Seth Weinberg for providing MATLAB code for the ORd cable, which served as the starting point for our code for the ToR-ORd and mKC cables. We thank Dr. Justin Morrisette-McCalmon for providing the raw experimental data which we analyzed and summarized in **Table 1**.

## REFERENCES

1. Basser PJ & Roth BJ (2000). New Currents in Electrical Stimulation of Excitable Tissues1. Annu Rev Biomed Eng 2, 377–397.

2. Boyle PM, Franceschi WH, Constantin M, Hawks C, Desplantez T, Trayanova NA & Vigmond EJ (2019). New insights on the cardiac safety factor: Unraveling the relationship between conduction velocity and robustness of propagation. J Mol Cell Cardiol 128, 117–128.

3. Boyle PM & Vigmond EJ (2010). An Intuitive Safety Factor for Cardiac Propagation. Biophys J 98, L57–L59.

4. Cheng Y-C et al. (2023). Combined Treatment of Human Induced Pluripotent Stem Cell– Derived Cardiomyocytes and Endothelial Cells Regenerate the Infarcted Heart in Mice and Non-Human Primates. Circulation 148, 1395–1409.

5. Coates S & Thwaites B (2000). The Strength-Duration Curve and Its Importance in Pacing Efficiency: A Study of 325 Pacing Leads in 229 Patients. Pacing Clin Electrophysiol 23, 1273–1277.

6. Colatsky T, Fermini B, Gintant G, Pierson JB, Sager P, Sekino Y, Strauss DG & Stockbridge N (2016). The Comprehensive in Vitro Proarrhythmia Assay (CiPA) initiative — Update on progress. J Pharmacol Toxicol Methods 81, 15–20.

7. Crestani T, Steichen C, Neri E, Rodrigues M, Fonseca-Alaniz MH, Ormrod B, Holt MR, Pandey P, Harding S, Ehler E & Krieger JE (2020). Electrical stimulation applied during differentiation drives the hiPSC-CMs towards a mature cardiac conduction-like cells. Biochem Biophys Res Commun 533, 376–382.

8. Davidenko JM, Levi RJ, Maid G, Elizari MV & Rosenbaum MB (1990). Rate dependence and supernormality in excitability of guinea pig papillary muscle. Am J Physiol-Heart Circ Physiol 259, H290–H299.

9. Du DTM, Hellen N, Kane C & Terracciano CMN (2015). Action Potential Morphology of Human Induced Pluripotent Stem Cell-Derived Cardiomyocytes Does Not Predict Cardiac Chamber Specificity and Is Dependent on Cell Density. Biophys J 108, 1–4.

10. Fermini B et al. (2016). A New Perspective in the Field of Cardiac Safety Testing through the Comprehensive In Vitro Proarrhythmia Assay Paradigm. J Biomol Screen 21, 1–11.

11. Geddes LA (2004). Accuracy limitations of chronaxie values. IEEE Trans Biomed Eng 51, 176–181.

12. Geddes LA & Bourland JD (1985). Tissue stimulation: Theoretical considerations and practical applications. Med Biol Eng Comput 23, 131–137.

13. Glukhov AV, Fedorov VV, Lou Q, Ravikumar VK, Kalish PW, Schuessler RB, Moazami N & Efimov IR (2010). Transmural Dispersion of Repolarization in Failing and Nonfailing Human Ventricle. Circ Res 106, 981–991.

14. Gong JQX & Sobie EA (2018). Population-based mechanistic modeling allows for quantitative predictions of drug responses across cell types. Npj Syst Biol Appl 4, 1–11.

15. Goversen B, van der Heyden MAG, van Veen TAB & de Boer TP (2018). The immature electrophysiological phenotype of iPSC-CMs still hampers *in vitro* drug screening: Special focus on IK1. Pharmacol Ther 183, 127–136.

16. Gray RA, Mashburn DN, Sidorov VY & Wikswo JP (2013). Quantification of Transmembrane Currents during Action Potential Propagation in the Heart. Biophys J 104, 268–278.

17. Gunawan MG, Sangha SS, Shafaattalab S, Lin E, Heims-Waldron DA, Bezzerides VJ, Laksman Z & Tibbits GF (2021). Drug screening platform using human induced pluripotent stem cell-derived atrial cardiomyocytes and optical mapping. STEM CELLS Transl Med 10, 68–82.

18. Gutstein DE, Morley GE, Tamaddon H, Vaidya D, Schneider MD, Chen J, Chien KR, Stuhlmann H & Fishman GI (2001). Conduction Slowing and Sudden Arrhythmic Death in Mice With Cardiac-Restricted Inactivation of Connexin43. Circ Res 88, 333–339.

19. Hawthorne RN, Blazeski A, Lowenthal J, Kannan S, Teuben R, DiSilvestre D, Morrissette-McAlmon J, Saffitz JE, Boheler KR, James CA, Chelko SP, Tomaselli G & Tung L (2021). Altered Electrical, Biomolecular, and Immunologic Phenotypes in a Novel Patient-Derived Stem Cell Model of Desmoglein-2 Mutant ARVC. J Clin Med 10, 3061.

20. He J-Q, Ma Y, Lee Y, Thomson JA & Kamp TJ (2003). Human Embryonic Stem Cells Develop Into Multiple Types of Cardiac Myocytes. Circ Res 93, 32–39.

21. Hodgkin AL (1954). A note on conduction velocity. J Physiol 125, 221–224.

22. Hoffman BF, Cranefield PF & Johnston FD (1960). Electrophysiology of the heart /. McGraw-Hill, New York.

23. Hund TJ & Rudy Y (2000). Determinants of Excitability in Cardiac Myocytes: Mechanistic Investigation of Memory Effect. Biophys J 79, 3095–3104.

24. Hwang HS, Kryshtal DO, Feaster TK, Sánchez-Freire V, Zhang J, Kamp TJ, Hong CC, Wu JC & Knollmann BC (2015). Comparable calcium handling of human iPSC-derived cardiomyocytes generated by multiple laboratories. J Mol Cell Cardiol 85, 79–88.

25. Hyde ER, Behar JM, Claridge S, Jackson T, Lee AWC, Remme EW, Sohal M, Plank G, Razavi R, Rinaldi CA & Niederer SA (2015). Beneficial Effect on Cardiac Resynchronization From Left Ventricular Endocardial Pacing Is Mediated by Early Access to High Conduction Velocity Tissue. Circ Arrhythm Electrophysiol 8, 1164–1172.

26. Jastrzębski M, Moskal P, Bednarek A, Kiełbasa G, Vijayaraman P & Czarnecka D (2019). His bundle has a shorter chronaxie than does the adjacent ventricular myocardium: Implications for pacemaker programming. Heart Rhythm 16, 1808–1816.

27. Kay GN, Mulholland DH, Epstein AE & Plumb VJ (1990). Effect of pacing rate on the human atrial strength-duration curve. J Am Coll Cardiol 15, 1618–1623.

28. Kernik DC, Morotti S, Wu H, Garg P, Duff HJ, Kurokawa J, Jalife J, Wu JC, Grandi E & Clancy CE (2019). A computational model of induced pluripotent stem-cell derived cardiomyocytes incorporating experimental variability from multiple data sources. J Physiol 597, 4533–4564.

29. Koivumäki JT, Naumenko N, Tuomainen T, Takalo J, Oksanen M, Puttonen KA, Lehtonen Š, Kuusisto J, Laakso M, Koistinaho J & Tavi P (2018). Structural Immaturity of Human iPSC-Derived Cardiomyocytes: In Silico Investigation of Effects on Function and Disease Modeling. Front Physiol; DOI: 10.3389/fphys.2018.00080.

30. Lapicque L (1907). Recherches quantitatives sur l’excitation electrique des nerfs traitee comme une polarization. J Physiol Pathol Générale 9, 620–635.

31. Lemoine MD, Mannhardt I, Breckwoldt K, Prondzynski M, Flenner F, Ulmer B, Hirt MN, Neuber C, Horváth A, Kloth B, Reichenspurner H, Willems S, Hansen A, Eschenhagen T & Christ T (2017). Human iPSC-derived cardiomyocytes cultured in 3D engineered heart tissue show physiological upstroke velocity and sodium current density. Sci Rep 7, 5464.

32. Li W, Han JL & Entcheva E (2020). Syncytium cell growth increases Kir2.1 contribution in human iPSC-cardiomyocytes. Am J Physiol-Heart Circ Physiol 319, H1112–H1122.

33. Lin J, Abraham A, George SA, Greer-Short A, Blair GA, Moreno A, Alber BR, Kay MW & Poelzing S (2022). Ephaptic Coupling Is a Mechanism of Conduction Reserve During Reduced Gap Junction Coupling. Front Physiol; DOI: 10.3389/fphys.2022.848019.

34. Ma J, Guo L, Fiene SJ, Anson BD, Thomson JA, Kamp TJ, Kolaja KL, Swanson BJ & January CT (2011). High purity human-induced pluripotent stem cell-derived cardiomyocytes: electrophysiological properties of action potentials and ionic currents. Am J Physiol-Heart Circ Physiol 301, H2006–H2017.

35. Miyaoka Y, Chan AH, Judge LM, Yoo J, Huang M, Nguyen TD, Lizarraga PP, So P-L & Conklin BR (2014). Isolation of single-base genome-edited human iPS cells without antibiotic selection. Nat Methods 11, 291–293.

36. Morotti S & Grandi E (2017). Logistic regression analysis of populations of electrophysiological models to assess proarrythmic risk. MethodsX 4, 25–34.

37. Muszkiewicz A, Britton OJ, Gemmell P, Passini E, Sánchez C, Zhou X, Carusi A, Quinn TA, Burrage K, Bueno-Orovio A & Rodriguez B (2016). Variability in cardiac electrophysiology: Using experimentally-calibrated populations of models to move beyond the single virtual physiological human paradigm. Prog Biophys Mol Biol 120, 115–127.

38. Ni H, Morotti S & Grandi E (2018). A Heart for Diversity: Simulating Variability in Cardiac Arrhythmia Research. Front Physiol; DOI: 10.3389/fphys.2018.00958.

39. Paci M, Hyttinen J, Aalto-Setälä K & Severi S (2013). Computational Models of Ventricular- and Atrial-Like Human Induced Pluripotent Stem Cell Derived Cardiomyocytes. Ann Biomed Eng 41, 2334–2348.

40. Paci M, Passini E, Klimas A, Severi S, Hyttinen J, Rodriguez B & Entcheva E (2020). All-Optical Electrophysiology Refines Populations of In Silico Human iPSC-CMs for Drug Evaluation. Biophys J 118, 2596–2611.

41. Plonsey R & Barr RC (2007). Bioelectricity: A Quantitative Approach. Springer Science & Business Media.

42. Pullinger TK & Sobie EA (2024). Cell-to-cell heterogeneity in ion channel conductance impacts substrate vulnerability to arrhythmia. Am J Physiol-Heart Circ Physiol; DOI: 10.1152/ajpheart.00645.2023.

43. Romagnuolo R et al. (2019). Human Embryonic Stem Cell-Derived Cardiomyocytes Regenerate the Infarcted Pig Heart but Induce Ventricular Tachyarrhythmias. Stem Cell Rep 12, 967–981.

44. Ronaldson-Bouchard K, Ma SP, Yeager K, Chen T, Song L, Sirabella D, Morikawa K, Teles D, Yazawa M & Vunjak-Novakovic G (2018). Advanced maturation of human cardiac tissue grown from pluripotent stem cells. Nature 556, 239–243.

45. Sarkar AX & Sobie EA (2011). Quantification of repolarization reserve to understand interpatient variability in the response to proarrhythmic drugs: A computational analysis. Heart Rhythm 8, 1749–1755.

46. Sayegh MN, Fernandez N & Cho HC (2019). Strength-duration relationship as a tool to prioritize cardiac tissue properties that govern electrical excitability. Am J Physiol-Heart Circ Physiol 317, H13–H25.

47. Seibertz F et al. (2023). Electrophysiological and calcium-handling development during long-term culture of human-induced pluripotent stem cell-derived cardiomyocytes. Basic Res Cardiol 118, 14.

48. Shah RR & Hondeghem LM (2005). Refining detection of drug-induced proarrhythmia: QT interval and TRIaD. Heart Rhythm 2, 758–772.

49. Sharma V & Tung L (2002). Spatial heterogeneity of transmembrane potential responses of single guinea-pig cardiac cells during electric field stimulation. J Physiol 542, 477– 492.

50. Shaw RM & Rudy Y (1997). Ionic Mechanisms of Propagation in Cardiac Tissue. Circ Res 81, 727–741.

51. Shiba Y, Gomibuchi T, Seto T, Wada Y, Ichimura H, Tanaka Y, Ogasawara T, Okada K, Shiba N, Sakamoto K, Ido D, Shiina T, Ohkura M, Nakai J, Uno N, Kazuki Y, Oshimura M, Minami I & Ikeda U (2016). Allogeneic transplantation of iPS cell-derived cardiomyocytes regenerates primate hearts. Nature 538, 388–391.

52. Sobie EA (2009). Parameter Sensitivity Analysis in Electrophysiological Models Using Multivariable Regression. Biophys J 96, 1264–1274.

53. Sobie EA, Susil RC & Tung L (1997). A generalized activating function for predicting virtual electrodes in cardiac tissue. Biophys J 73, 1410–1423.

54. Sottas V, Wahl C-M, Trache MC, Bartolf-Kopp M, Cambridge S, Hecker M & Ullrich ND (2018). Improving electrical properties of iPSC-cardiomyocytes by enhancing Cx43 expression. J Mol Cell Cardiol 120, 31–41.

55. Sridharan D, Pracha N, Rana SJ, Ahmed S, Dewani AJ, Alvi SB, Mergaye M, Ahmed U & Khan M (2023). Preclinical Large Animal Porcine Models for Cardiac Regeneration and Its Clinical Translation: Role of hiPSC-Derived Cardiomyocytes. Cells 12, 1090.

56. Starmer CF, Grant AO & Strauss HC (1984). Mechanisms of use-dependent block of sodium channels in excitable membranes by local anesthetics. Biophys J 46, 15–27.

57. Tomek J, Bueno-Orovio A, Passini E, Zhou X, Minchole A, Britton O, Bartolucci C, Severi S, Shrier A, Virag L, Varro A & Rodriguez B (2019). Development, calibration, and validation of a novel human ventricular myocyte model in health, disease, and drug block ed. Faraldo-Gómez JD, Barkai N, Hund T & Maleckar M. eLife 8, e48890.

58. Tung L & Borderies JR (1992). Analysis of electric field stimulation of single cardiac muscle cells. Biophys J 63, 371–386.

59. Tung L & Kléber AG (2000). Virtual sources associated with linear and curved strands of cardiac cells. Am J Physiol-Heart Circ Physiol 279, H1579–H1590.

60. Tung L, Sliz N & Mulligan MR (1991). Influence of electrical axis of stimulation on excitation of cardiac muscle cells. Circ Res 69, 722–730.

61. Vaidyanathan R, Markandeya YS, Kamp TJ, Makielski JC, January CT & Eckhardt LL (2016). IK1-enhanced human-induced pluripotent stem cell-derived cardiomyocytes: an improved cardiomyocyte model to investigate inherited arrhythmia syndromes. Am J Physiol-Heart Circ Physiol 310, H1611–H1621.

62. Weinberg SH (2017). Ephaptic coupling rescues conduction failure in weakly coupled cardiac tissue with voltage-gated gap junctions. Chaos Interdiscip J Nonlinear Sci 27, 093908.

63. Weiss G (1901). Sur la possibilite de rendre comparables entre eux les appareils servant a l’excitation electrique. Arch Ital Biol 35, 413–445.

64. Yang J, Daily N, Pullinger TK, Wakatsuki T & Sobie EA (2024). Creating cell-specific computational models of stem cell-derived cardiomyocytes using optical experiments. 2024.01.07.574577. Available at: https://www.biorxiv.org/content/10.1101/2024.01.07.574577v1 [Accessed March 1, 2024].

65. Zhang J, Wilson GF, Soerens AG, Koonce CH, Yu J, Palecek SP, Thomson JA & Kamp TJ (2009). Functional Cardiomyocytes Derived From Human Induced Pluripotent Stem Cells. Circ Res 104, e30–e41.

